# The microtubule-binding protein EML3 is required for mammalian embryonic growth and cerebral cortical development; *Eml3* null mice are a model of cobblestone brain malformation

**DOI:** 10.1101/2025.04.06.647459

**Authors:** Isabelle Carrier, Eduardo Diez, Valerio E.C. Piscopo, Susanne Bechstedt, Hans van Bokhoven, Myriam Srour, Albert Berghuis, Stefano Stifani, Yojiro Yamanaka, Roderick R. McInnes

## Abstract

The cerebral cortex is a multi-layered structure generated through the migration of neural precursors from their birthplace in the ventricular zone to their destination within the cortical plate. Neuronal migration defects are responsible for many human pathologies collectively called neuronal migration disorders, which include subcortical band heterotopia and cobblestone brain (COB) malformation. One example of a protein involved in a neuronal migration disorder is the echinoderm microtubule-associated protein-like 1 (EML1) protein, one of six members of the mammalian EML family. Absence of EML1 protein results in subcortical band heterotopia in mice and humans. Here, we report that absence of the paralogous protein EML3 leads to delayed embryonic development and small size, and a COB-like phenotype with neuronal ectopias in the dorsal telencephalon. We found that EML3 is expressed in the neuroepithelium and meningeal mesenchyme when those tissues participate in pial basement membrane (PBM) formation. Transmission electron microscopy demonstrated that the extracellular matrix of the PBM is structurally abnormal in *Eml3* null mice when the first radially migrating neurons arrive. The reduced structural integrity of the PBM leads to focal over-migration of neurons into the subarachnoid space. These findings strengthen the link between the EML protein family and cortical neuronal migration defects by identifying *Eml3* as the first EML family member whose absence leads to over-migration of neuroblasts. Moreover, we report the first COB-like phenotype with PBM structural defects when a single microtubule-associated protein is deleted.

## INTRODUCTION

The cerebral cortex is generated through a series of steps that include the migration of neural precursors (neuroblasts) from their birthplace in the ventricular zone to their destination in the cortical plate (Gupta et al 2002, Noctor et al 2008). Neural precursors arise from progenitor cells called radial glia that are themselves differentiated from neuroepithelial cells beginning at embryonic day 10.5 (E10.5) in the mouse (Gotz & Huttner 2005, Noctor et al 2008). Both neuroepithelial cells and radial glia are polarized cells that span the entire developing cerebral cortex (Gotz & Huttner 2005, Noctor et al 2008). Neuroepithelial cells have a strictly proliferative (symmetric) cell division mode that yields two identical daughter cells (Kriegstein & Alvarez-Buylla 2009, Taverna & Huttner 2010). Radial glia, in contrast, switch to a neurogenic (asymmetric) mode of cell division that yields one radial glia and a neuroblast committed to becoming a neuron during neurogenesis (Noctor et al 2008, Taverna & Huttner 2010). The mammalian cortex develops in an inside-out manner, with radially migrating neuroblasts traversing earlier-formed layers to reach the cortical plate’s outer surface (Noctor et al 2004, Valiente & Marin 2010). Each layer emerges with precise timing, regulated by defined molecular and transcriptional cues (Bayatti et al 2008, Bedogni et al 2010, Hoerder-Suabedissen & Molnar 2015). Radial glial fibers guide the neuroblasts along the radial axis. Thus, radial glia serve as both neural progenitors and scaffolds for migrating neurons (Anthony et al 2004, Gotz & Huttner 2005). In mice, neuroblast radial migration starts just before E11.5 and is completed by E16.5 (Kriegstein et al 2006, Miyata et al 2004).

There are physical boundaries as well as molecular stop signals that define when a migrating neuroblast has reached its destination (Kawauchi 2012). The pial basement membrane (PBM) serves both as a physical barrier and as a source of molecular cues (Siegenthaler & Pleasure 2011). The PBM is made up of an extracellular matrix (ECM) and the basal endfeet of neuroepithelial cells or radial glia depending on the embryonic developmental stage (Belvindrah et al 2011, Gotz & Huttner 2005). The ECM components are secreted from the neural precursors as well as from the overlaying mesenchymal cells (Franco & Muller 2011). Several ECM components have self-assembling properties and also interact with cell surface receptors on the basal endfeet of neural precursors, thus anchoring and stabilizing the ECM (Chiquet-Ehrismann & Tucker 2011, Yurchenco 2015). Once the ECM is synthesized and deposited, it undergoes constant remodeling (Bonnans et al 2014, Maeda 2015).

Cortical neuronal migration defects cause alterations in the structure of the neocortex, and are frequently associated with severe epilepsy (Barkovich et al 2012, Guerrini & Dobyns 2014). These disorders include lissencephaly, pachygyria, polymicrogyria, periventricular nodular heterotopia, subcortical band heterotopia, and cobblestone brain malformation (Barkovich et al 2012, Guerrini & Filippi 2005). Cobblestone brain malformation (COB) is a neuronal over-migration condition characterized by an irregular brain surface with nodular protrusions into the overlaying subarachnoid space (Dobyns & Truwit 1995, Guerrini & Dobyns 2014). In contrast, subcortical band heterotopia is due to a failure of neurons to migrate to the cerebral cortex, instead forming a band of gray matter underneath the cortical surface (neuronal under-migration) (Guerrini & Dobyns 2014, Guerrini & Marini 2006). The Echinoderm Microtubule-associated Protein-like 1 (*EML1*) gene has been associated with subcortical band heterotopia (Kielar et al 2014, Zaidi et al 2024). Abnormal mitotic spindle orientation has been observed in *Eml1*-null progenitors and the ensuing delamination of apical progenitors has been proposed as leading to the ectopias (Bizzotto et al 2017, Kielar et al 2014). Although EML1 is a member of a family of proteins that includes six members in both mice and humans (Fry et al 2016), EML1 is thus far the only member that has been involved in cortical malformations.

COB (formerly referred to as lissencephaly type 2) is characterized by over-migrating neuroblasts passing through breaches in the PBM into the subarachnoid space (Barkovich et al 2015, Barkovich et al 2012, Guerrini & Dobyns 2014, Leventer et al 2008). There are two alternative pathogenic mechanisms that can result in COB. The most common mechanism involves a weak or disrupted PBM that fails to provide a physical boundary or chemical stop cues to migrating neurons (Radmanesh et al 2013, van Reeuwijk et al 2005b). In the alternative mechanism, migrating neurons fail to detect or interpret stop cues and ultimately tear through an intact and functional PBM (Moers et al 2008). COB is associated with mutations in the *DAG1* gene, encoding for the alpha-dystroglycan protein (Frost et al 2010, Geis et al 2013, Riemersma et al 2015), and those involved in its glycosylation, i.e. glycosyltransferases *POMT1*, *POMT2*, *FKTN*, *FKRP*, *POMGNT1*, and *LARGE* (Beltran-Valero de Bernabe et al 2002, Godfrey et al 2007, Mercuri et al 2009, van Reeuwijk et al 2005a, Willer et al 2014). These mutations disrupt radial glial basal endfoot binding to the ECM, destabilizing the PBM (Moore et al 2002). Mutations in ECM components like laminin gamma-1 can also contribute to PBM instability (Halfter et al 2002). Since the alpha-dystroglycan and laminin gamma-1 proteins are also present in basement membranes other than the PBM, COB is typically observed as one of the phenotypes of a syndrome (Godfrey et al 2007, Mercuri et al 2009, Willem et al 2002). However, COB can also present as an isolated condition, with about 30% of COB cases exhibiting atypical non-syndromic forms with no identified mutations, underscoring the disorder’s significant genetic and clinical heterogeneity (Devisme et al 2012, Radmanesh et al 2013).

The EML proteins contain a tandem atypical propeller in EMLs (TAPE) domain containing WD40 repeats that fold into beta-propellers (Richards et al 2014). The C-terminal TAPE domain in EML1 and EML2 has been shown to bind soluble tubulins (Richards et al 2014). The N-terminus of EML proteins harbors a coiled-coil which has been demonstrated in EML2 and EML4 to mediate protein trimerization (Richards et al 2015). Additionally, in EML1 and EML4, the N-terminal region was shown to be required for association with polymerized microtubules (Pollmann et al 2006, Richards et al 2015). Although EML proteins have been proposed to modulate microtubule stability (Eichenmuller et al 2002, Fry et al 2016, Houtman et al 2007, Pollmann et al 2006), further research is needed to understand the precise functions and contributions of each member of the EML protein family in cellular processes and specifically in neuronal development.

Previous work has shown that when *EML3* is knocked down in cultured cells, chromosomes are misaligned during metaphase and cells are delayed in the mitotic phase of the cell cycle (Tegha-Dunghu et al 2008). It was later shown that EML3 is required for recruitment of the augmin protein and gamma-tubulin ring complexes to existing microtubules for microtubule-based microtubule nucleation, a process that enables mitotic spindles to capture metaphase chromosome kinetochores in a timely manner (Luo et al 2019). These studies define EML3 as a modulator of spindle assembly during cell division. Given the role for EML3 in mitotic spindle formation in cultured cells and given the role for EML1 in controlling spindle orientation in neural progenitors we hypothesized that the *Eml3* gene may also be required for cortical development.

To examine the possibility that *Eml3* is required during development we generated *Eml3* null mice. We report that *Eml3* null embryos are developmentally delayed, die perinatally, and have focal neuronal ectopias with neuroblasts that migrate into the overlaying subarachnoid space through a defective PBM. *Eml3* null mice are a model for cobblestone brain malformation with a defect in ECM remodeling.

## RESULTS

### Generation of *Eml3* null mice

To characterize the developmental processes requiring the EML3 protein, the mouse *Eml3* gene was targeted. As described in Materials and Methods, the following mice were generated: *Eml3^tm1e(EUCOMM)Wtsi^* or *Eml3*^gt^ (gene trap allele) and *Eml3^em1.2Mci^* or *Eml3*^-^ (knockout allele with exons 11-19 deleted) (Supp. Fig. 1). Both mouse lines were verified to represent *Eml3* null alleles. The reduction of EML3 protein levels in heterozygotes (*Eml3^wt/gt^* and *Eml3^+/-^*) and its elimination from homozygous embryos (*Eml3^gt/gt^*and *Eml3^-/-^*) was confirmed by immunoblotting (Supp. Fig. 1F and 1G). Mice were backcrossed to C57BL/6J background for at least two generations prior to conducting experiments. Experimental mice were produced by intercrossing heterozygotes. All experiments were performed using both male and female littermates and all data shown is pooled from both sexes after verification that there were no sex-dependent differences.

Three main phenotypes were observed in *Eml3* null mice: embryonic development delay, brain defect, and perinatal lethality (Table 1). All phenotypes are recessive, heterozygous mice (*Eml3^+/-^*) being indistinguishable from their *Eml3^+/+^* littermates.

**Table 1.** Phenotypes observed in Eml3 null mice.

| Phenotype | Sub-phenotype | Age when observed <sup>a</sup> | Penetrance of phenotype vs *Eml3* genotype |  |  |
| --- | --- | --- | --- | --- | --- |
|  |  |  | *Eml3*<sup>+/+</sup> | *Eml3*<sup>+/-</sup> | *Eml3*<sup>-/-</sup> |
| **Embryo development delay** | Small embryo and placenta size <sup>b</sup> | E8.5-<u>E18.5</u> | 2/41 (4.9%) | 1/75 (1.3%) | 23/23 (100%) |
|  | Fewer somite pairs <sup>c</sup> | E8.5-<u>E12.5</u> | 3/25 (12%) | 4/51 (8%) | 11/20 (55%) |
|  | Delayed neurogenesis <sup>d</sup> | E10.5-<u>E11.5</u> | 0/10 (0%) | 0/1 (0%) | 8/11 (72.7%) |
|  | Thin corpus callosum <sup>e</sup> | E18.5 | 0/8 (0%) | 0/1 (0%) | 10/10 (100%) |
| **Brain defect** | Focal neuronal ectopias | E11.5-<u>E18.5</u> | 0/10 (0%) | n.d. | 10/14 (71.4%) |
| **Perinatal lethality** | Failure to inflate lungs <sup>f</sup> | E18.5 | 5/24 (20.8%) | 5/33 (15.2%) | 6/10 (60%) |
|  | Survival up to weaning age <sup>g</sup> | PN18-24 | n.d. | 355 Het vs 191 WT (~7% Het lethality) | 11 KO vs 191 WT (~94% KO lethality) |

<sup>a</sup> Embryonic (E) or postnatal (PN) ages at which the phenotype was observed. When a range of ages were studied, the penetrance data shown in the table is from the underlined age.
<sup>b</sup> Embryo sizes were measured as area of the section corresponding to the sagittally bisected E8.5-E10.5 embryos or as weights for E11.5-E18.5 embryos. E18.5 weight analyses shown, with 15% lighter weight vs average of Heterozygote littermates as threshold for small size. 100% penetrance for low weight of *Eml3* null embryos was observed at E12.5-E18.5. *Eml3* null placentas are small at E10.5-E18.5.
<sup>c</sup> Somite pairs were counted under a dissection microscope for E8.5-E12.5 embryos. E12.5 data shown, with a minimum difference of 1.5 somite pairs vs the average for heterozygous littermates as the threshold for delay.
<sup>d</sup> Delayed neurogenesis was determined by staining E10.5-E11.5 frontal cortices with markers for neuroepithelial progenitors (PAX6 and SOX1), intermediate neuronal progenitors (TBR2), and postmitotic projection neurons (TBR1). E11.5 data shown as number of embryos with at least a 20% lower percentage of TBR2-positive cells in their frontal cortex as compared to a control littermate.
<sup>e</sup> Corpus callosum (CC) thicknesses were measured in Nissl-stained coronal sections of E18.5 whole brains. Pairwise comparisons among littermates determined that all *Eml3* null embryos had a disproportionately thinner CC than their littermates even when normalizing for the overall smaller size of the embryos.
<sup>f</sup> Air breathing and lung floating assays were performed on E18.5 embryos collected by caesarean section. The pups were observed and filmed for 90 minutes. At the end of the observation period all mice were euthanized, and the lungs were removed and placed in PBS for the lung floating assay.
<sup>g</sup> We estimate the number of missing mice at weaning age by comparing the number of *Eml3* null and heterozygous mice to the number of weaned wild type mice. Given that we analyze litters from heterozygote intercrosses and that we expect a ratio of 1WT:2Het:1KO, and that we obtained a total of 191 WT mice, we were expected to recover 382 Het and 191 KO mice.

### Perinatal lethality in *Eml3* null mice

To determine the phenotype of *Eml3* null mice, we intercrossed heterozygotes (*Eml3*^+/-^) and characterized the offspring that survived until weaning age (approximately 21 days old). Out of 557 weaned mice only 11 *Eml3*^-/-^ pups were obtained (Tables 1 and 2). *Eml3*^-/-^ mice, therefore, represent 2% of all weaned mice, in contrast with the expected 25% if viable. Notably, most of the rare *Eml3* null survivors were runted with 32% average lower body weight than littermate controls and 25% had hydrocephalus. The lethality phenotype is recessive since heterozygotes (*Eml3*^+/-^) are found approximately in the expected proportion of 2:1 versus wild type mice (*Eml3*^+/+^). We conclude that a majority of *Eml3* null mice die before weaning age and the phenotype is recessive.

**Table 2.**
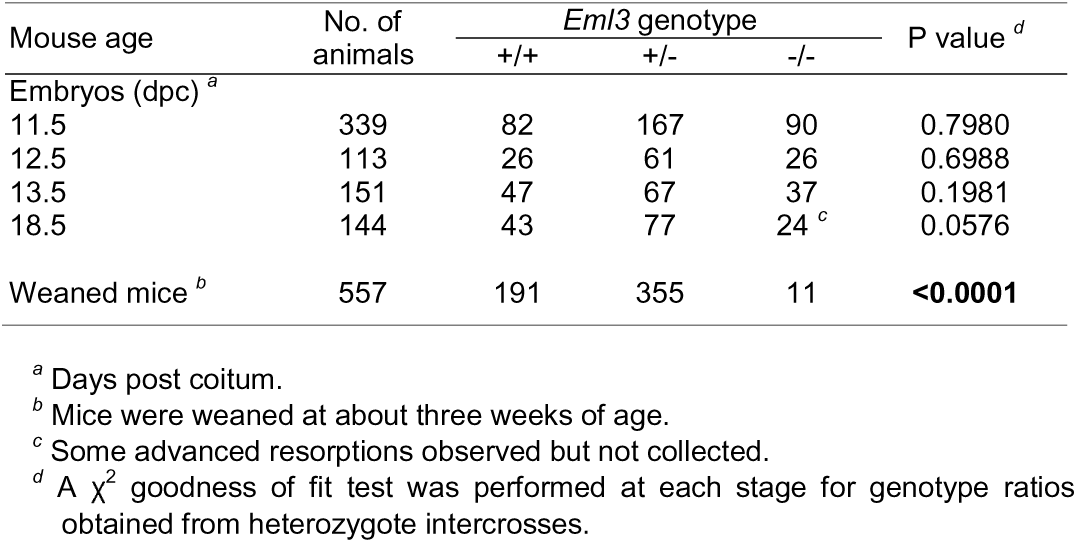
Lethality in Eml3 null mice.

To define the timepoint of lethality, timed heterozygote intercross matings were set up and genotype ratios were determined at several stages of embryonic development (Table 2). Mendelian ratios were obtained at all stages studied according to chi-square analyses. We conclude that *Eml3* null mice die mostly during or after birth. Dead *Eml3* null pups were observed within 48 hours following birth. To gain insight into the underlying cause of perinatal death, we closely monitored the fate of litters obtained via C-section at E18.5. The observation period lasted for 1.5 hours after surgical delivery. 6/10 (60%) *Eml3* null neonates and 10/57 (18%) control littermates showed severe respiratory distress and died within 30-40 min after C-section (Table 1). These neonates had strong respiratory reflexes but remained cyanotic and died of acute respiratory failure. Immediately following the observation period, the lungs were dissected from the neonates and immersed in saline. Lungs from neonates that had not survived sank in the saline since they had not inflated with air. Failure to inflate lungs can be a consequence of lung immaturity and is encountered in several knockout mouse models (Turgeon & Meloche 2009). Indeed, histological analysis of serial sections of E18.5 *Eml3* null embryos revealed immaturity of the lungs in a significant portion of the *Eml3* null embryos examined (3/4 *Eml3* null embryos had not reached saccular stage in lung development, a proportion higher than what was observed in control mice). No skeletal defect, nor structural defect in the diaphragm and intercostal muscles were observed in the thorax of *Eml3* null embryos. The analyses did not reveal any gross morphological anomaly in other internal organs. Overall, the neonate characterization data suggests that the postnatal lethality of *Eml3* null mice was at least in part due to respiratory distress due to lung immaturity.

### Global developmental delay in *Eml3* null embryos

While characterizing embryos and neonates, we noted that *Eml3* null mice were smaller than control littermates (Fig. 1A and data not shown). Except for their size, *Eml3* null embryos were morphologically normal throughout embryonic development (Fig. 1A). To determine if the lung immaturity and small size of *Eml3* null neonates are an indication of global developmental delay we characterized *Eml3* null embryos at different stages of development. Timed matings were established between *Eml3* heterozygotes and embryos were examined at specific ages, measured in days post coitum. At E7.5 no significant size difference was observed between *Eml3* null and control embryos (Fig. 1A). However, on embryonic days E8.5 to E18.5, they were 19-44% smaller in size (Fig. 1A). The placentas were also examined from E10.5 to E18.5 and found to be 20-25% smaller in size (Fig. 1B). In addition, delayed appearance of intermediate progenitors (Fig. 3A) and neuroblasts, delayed saccular stage of lungs at E18.5, and a thinner corpus callosum at E18.5 (Table 1) suggests that *Eml3* null embryos are delayed in development. Indeed, within a single litter, *Eml3* null embryos had fewer somite pairs when compared to control littermates at all stages analyzed (E8.5-E12.5; Fig. 1C). At E8.5, shortly after the onset of somitogenesis, there was on average one somite pair fewer in *Eml3* null embryos compared to control littermates (Fig. 1C). At E9.5, E10.5 and E11.5 a constant two somite pair mean reduction in the total number of somite pairs was observed. Interestingly, comparing the weight of *Eml3* null embryos to control embryos with the same number of somite pairs, i.e., stage-matched, revealed that *Eml3* null embryos were still smaller than control embryos (Fig. 1D). This suggests that, in addition to the developmental delay, the *Eml3* null embryos are growth restricted. We conclude that *Eml3* null mice are developmentally delayed starting at E8.5 and are smaller than stage-matched control littermates (growth restricted).

**Fig. 1.**
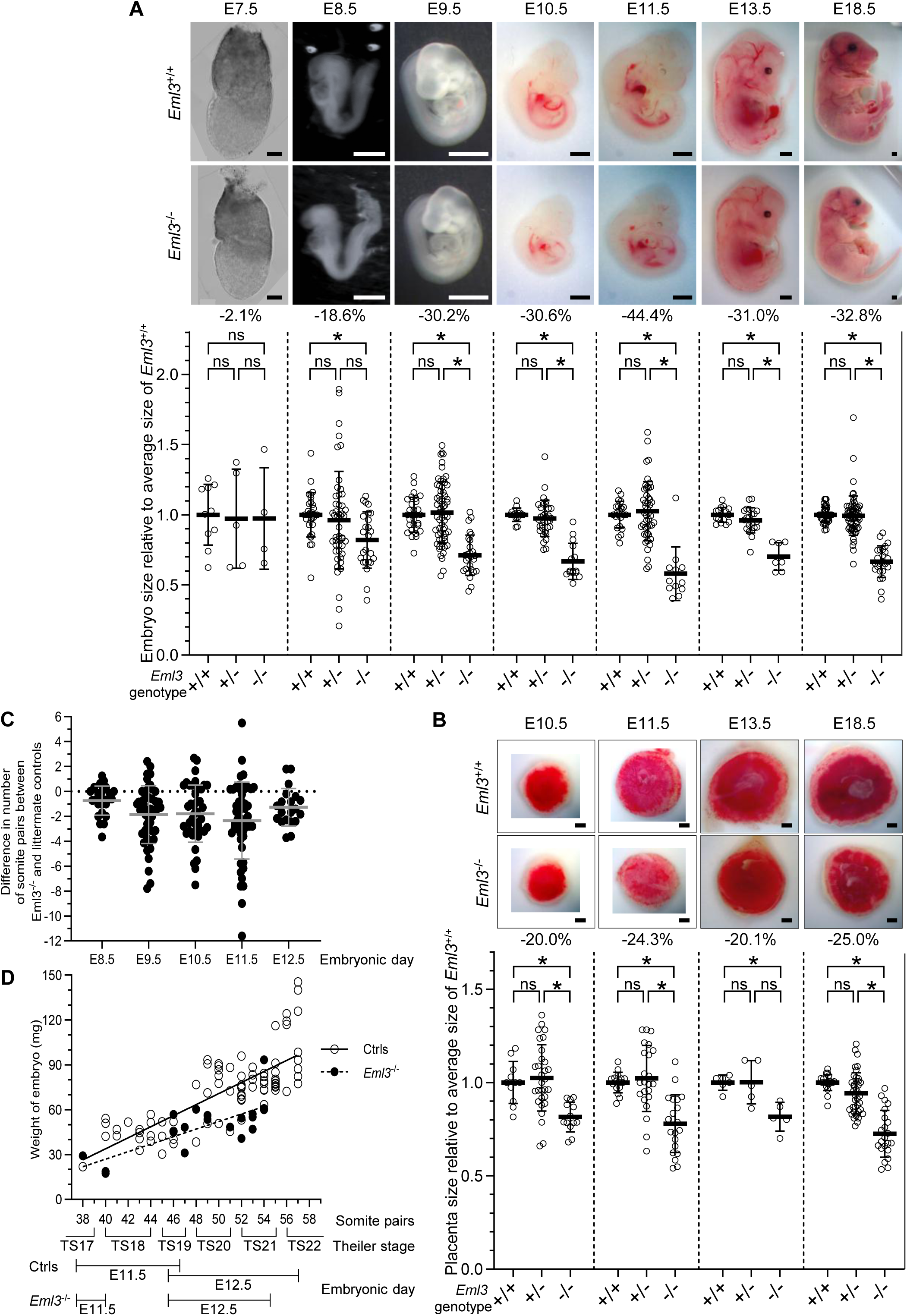
*Eml3^-^*^/-^ embryos are smaller than control littermates and are developmentally delayed. At each gestational age between E8.5 and E18.5, the *Eml3^-^*^/-^ embryos **(A)** and their placentas **(B)** (E10.5- E18.5) are significantly smaller than their *Eml3*^+/+^ and *Eml3*^+/-^ control littermates. The size was measured either by area (E7.5-E10.5 embryos; E10.5-E18.5 placentas) or weight (E11.5-E18.5 embryos); for each litter analyzed, the size of each embryo or placenta was measured relative to the average of the control embryos (*Eml3*^+/+^ and *Eml3*^+/-^) and then normalized to the average of the *Eml3*^+/+^ embryos. The average percent size difference is shown below each image (n = 5-32). Dunn’s pairwise comparison, * = p<0.05, ns = not significant. Scale bars, 100µm for E7.5, 500µm for E8.5, and 1mm for E9.5-E18.5. **(C)** At each gestational age between E8.5 and E12.5, the *Eml3*^-/-^ embryos have significantly fewer somite pairs than *Eml3*^+/+^ and *Eml3*^+/-^ control littermates; average difference in number of somite pairs ± SD shown (n = 20 - 50). **(D)** Weight of embryos was plotted against the number of somite pairs counted for *Eml3*^-/-^ and control embryos. Linear regression was used to fit the datapoints; *Eml3*^-/-^ embryos are significantly smaller (p<0.0001) than control embryos with the same number of somite pairs.

### Focal neuronal ectopias in *Eml3* null embryos

#### Regional lamination defects in Eml3 null mouse brains

To determine if the *Eml3* null brains have gross morphological defects we stained coronal and sagittal brain sections for the Nissl substance (cresyl violet stain) that highlights neuronal structure. In E18.5 *Eml3* null mouse embryos we observed ectopic clusters of cells within the marginal zone and the subarachnoid space of the dorsal telencephalon (Fig. 2A). These ectopias were observed in 10/14 *Eml3* null brains examined (71.4% penetrance) and absent from all 10 control *Eml3^+/+^* mice analyzed (Table 1). The cells in the ectopic clusters were immunoreactive when stained with antibodies to the pan-neuronal marker tubulin beta-3 (TUBB3; Fig. 2B), identifying the cells as neurons and establishing the ectopic clusters as focal neuronal ectopias (FNEs). Laminin immunostaining, a major component of the PBM, indicated that the PBM was fragmented at the FNEs (laminin speckles; Fig. 2B). Below each FNE the cortical architecture was disorganized suggesting that a cohort of neurons had over-migrated and traversed the pial basement membrane (Fig. 2A).

**Fig. 2.**
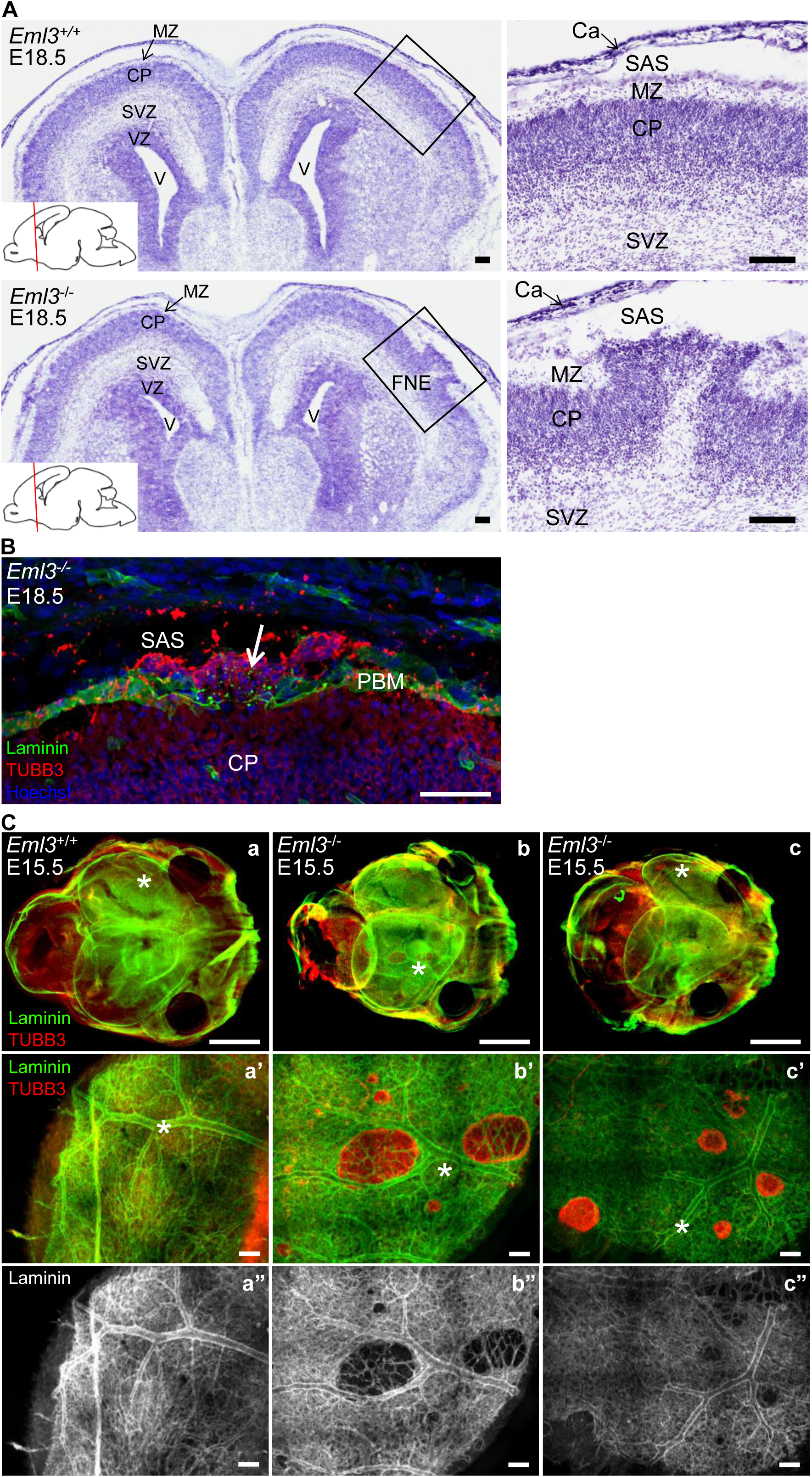
Focal neuronal ectopias (FNEs) in *Eml3^-/-^ embryos*. **(A)** Nissl (cresyl-violet) staining of coronal sections of forebrains from E18.5 *Eml3^+/+^* and *Eml3^-/-^* embryos. The *Eml3^+/+^* sections in the top panels show normal brain architecture with a close-up view in the panel on the right. An FNE is present in the section of the *Eml3^-/-^* cortex shown in the bottom panels. The close-up in the bottom-right panel shows over-migrating neurons (dark cresyl-violet staining) extending past the marginal zone and into the subarachnoid space. Ca, calvarium; CP, cortical plate; MZ, marginal zone; SAS, subarachnoid space; SVZ, subventricular zone; V, ventricle; VZ, ventricular zone. Scale bars, 100µm. **(B)** Immunolabelling of an E18.5 *Eml3^-/-^* mouse brain section for laminin (green) a major component of the extracellular matrix of the pial basement membrane, and for neuron-specific tubulin beta-3 (TUBB3, red). The pial basement membrane is disrupted at the FNEs, with neurons (TUBB3+) over-migrating into the subarachnoid space. Laminin speckles are observed (arrow). CP, cortical plate; PBM, pial basement membrane; SAS, subarachnoid space. Scale bar, 50µm. **(C)** Heads of E15.5 *Eml*3^+/+^ and *Eml3*^-/-^ embryos were immuno-stained and cleared before imaging with a fluorescence stereomicroscope **(a-c)** or with a confocal microscope **(a’-c”)**. Immunostaining was performed with antibodies against the ECM protein laminin (green in **a**-**c** and **a’**-**c’**, white in **a”**-**c”**) and a marker of terminally differentiated neurons, tubulin beta-3 (TUBB3, red). The asterisks point to the same area in the high magnification confocal microscope images as in the low magnification stereomicroscope images. Scale bars, 1mm **(a-c)** and 0.1mm **(a’-c”)**.

To analyze the spatial distribution of FNEs in developing brains, immunostained heads were cleared and imaged (Fig. 2C). Whole E15.5 heads were immunostained with laminin to highlight the pial basement membrane and tubulin beta-3 to highlight post-mitotic neuroblasts. In addition, *Eml3* null E18.5 brain serial sections were stained with cresyl violet (Fig. 2A). The number of FNEs in *Eml3* null brains varied from none to twelve, the FNEs varied in size and were randomly distributed rostrocaudally in both forebrain hemispheres (Fig. 2A and 2C). FNEs were restricted to the dorsal aspect of the telencephalon whereas medial cortical fields near the interhemispheric fissure and ventral areas were unaffected. No histological abnormalities were found in other brain regions.

#### Normal neurogenesis in Eml3 null dorsal telencephalon

To establish whether the focal neuronal ectopias observed in *Eml3* null mice are a consequence of disturbed neurogenesis, as determined for the subcortical band heterotopias present in *Eml1* null mice (Bizzotto et al 2017, Kielar et al 2014), we immunostained *Eml3* null and control forebrains with key markers of neurogenesis. E10.5 and E11.5 embryo cryosections were immunostained for TBR2 and TBR1 which are markers of intermediate progenitors and of post-mitotic neural precursors respectively. The appearance of both cell types defines the onset of neurogenesis. We calculated the percentage of TBR2 and TBR1 immunoreactive cells present in each forebrain cryosection at specific embryonic ages. The appearance of intermediate progenitors was delayed in *Eml3* null mice (TBR2 at E11.5, Fig. 3A). Analyses at additional embryonic ages and with TBR1 staining indicated that neurogenesis milestones were reached at a later age in *Eml3* null embryos (data not shown). Since *Eml3* null mice are developmentally delayed as determined by the number of somites at a specific age (Fig. 1C), we then plotted the average percentage of TBR2+ cells against the number of somite pairs for each embryo, allowing for a comparison of stage matched *Eml3* null and control embryos. TBR2+ cells appeared at the same stage and their prevalence increased at the same rate in *Eml3* null and control embryos (Fig. 3B and data not shown for TBR1). We tested two additional markers, SOX1 and PAX6, that are expressed sequentially during early neurogenesis. The transition from *Sox1* to *Pax6* expression in mice marks the progression from early neuroepithelium to more differentiated radial glial cells during early embryonic development, typically occurring around embryonic day E8.5-9.5. *Sox1* is expressed first, marking the neural plate, while *Pax6* is expressed subsequently, driving the transition to radial glia and neuronal differentiation (Suter et al 2009). No differences in SOX1 and PAX6 staining were observed between stage-matched *Eml3* null and control embryos (data not shown).

**Fig. 3.**
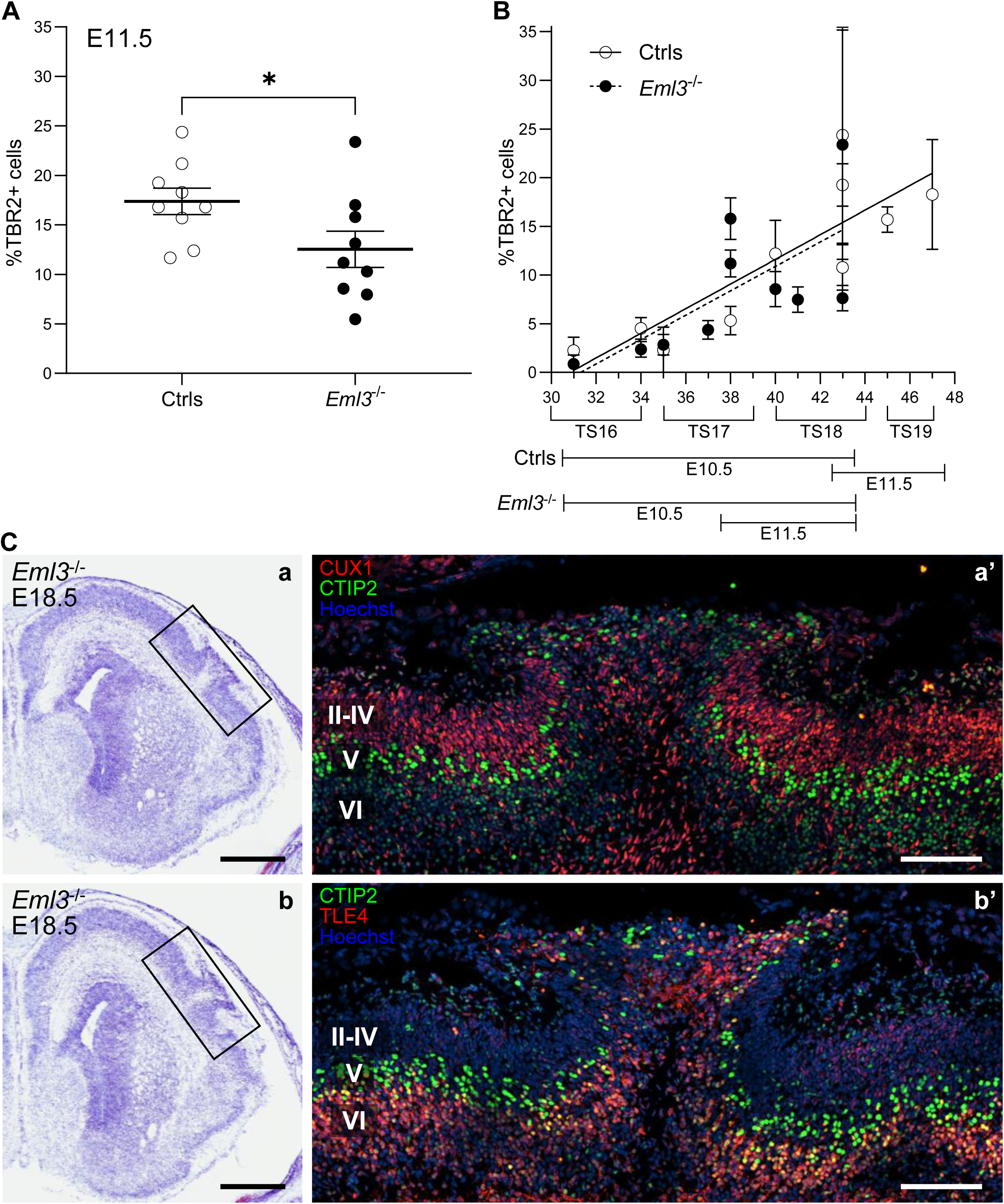
*Eml3*^-/-^ embryos are developmentally delayed *vs.* littermate controls but the onset and progression of neurogenesis is equal in somite-matched *Eml3^-/-^* and control forebrains; additionally, neurons of all cortical plate layers are present in *Eml3^-/-^* FNEs. **(A)** The onset and progression of neurogenesis is delayed in *Eml3*^-/-^ *vs.* littermate control forebrains. Forebrains from embryos collected at E11.5 were cryosectioned and processed for indirect immunofluorescence with markers for cell populations whose relative abundances indicate the onset and progression of neurogenesis. The percentage of TBR2-positive intermediate progenitors was determined. *Eml3^+/+^* and *Eml3^+/-^* data are pooled as controls (Ctrls) *vs. Eml3^-/-^* embryos. The total number of cells in the embryonic brain section was determined with the nuclear stain Hoechst. Each dot on the graph represents the mean percentage of cells positive for TBR2 from analysis of 2-4 sections per embryo. Error bars represent SEM. **(B)** The onset and progression of neurogenesis is equal in somite-matched *Eml3^-/-^* and control forebrains. Forebrains from embryos collected at E10.5 and E11.5 were cryosectioned and processed for indirect immunofluorescence with markers for cell populations whose relative abundances indicate the onset and progression of neurogenesis. The percentage of TBR2-positive intermediate progenitors was determined and is plotted against the number of somite pairs counted in each embryo. The total number of cells in the embryonic brain section was determined with the nuclear stain Hoechst. Each dot on the graph represents the mean percentage of cells positive for TBR2 in an individual embryo. Error bars represent SD from analysis of 2-4 sections per embryo. *Eml3^+/+^* and *Eml3^+/-^* data are pooled as controls (Ctrls) *vs. Eml3^-/-^* embryos. Shown below the graphs is the Theiler stage of development that corresponds to the number of somite pairs counted. Also indicated is the age of the embryos in days post-coitum. **(C)** Neurons of all cortical plate layers are present in *Eml3^-/-^* FNEs at E18.5. **(a, b)** Nissl (cresyl-violet) staining of serial coronal sections through a single FNE. **(a’, b’)** Immunolabeling of adjacent sections of the same FNE with markers of cortical plate layers. **(a’)** CUX1 immunostaining of late-born upper layer neurons (II-IV) and CTIP2 immunostaining of intermediate layer neurons (V). **(b’)** TLE4 immunostaining of early-born deeper layer neurons (VI) and CTIP2 immunostaining. Scale bars, 500µm **(a, b)** and 100µm **(a’, b’)**.

Thus, *Eml3* null mice have reached the same neurogenesis milestones as their WT counterparts when they have the same number of somites (Fig. 3B). Given that *Eml1* null radial glia have perturbed mitotic spindle angles, we measured the angle of mitotic spindles in *Eml3* null and control apical progenitors and saw no differences when the embryos were stage-matched (data not shown). We conclude that the onset and progression of neurogenesis is normal in *Eml3* null mice.

#### Ectopic neurons comprise neurons from all cortical layers

Having determined that the cortical defect in *Eml3* null mice is not associated with abnormal neurogenesis, we aimed to identify what other pathological mechanism is responsible for the *Eml3* null FNEs. Our first aim was to establish the onset of neuronal over-migration. Since neurons born at different developmental stages express different markers (Greig et al 2013, Noctor et al 2004), identifying the neurons present within the FNEs gives an estimate of the onset of neuronal over-migration. To determine the cellular composition of the ectopias observed at E18.5, we performed immunostaining with layer-specific markers: CUX1 for layers II–IV (Nieto et al 2004), CTIP2 for layer V (Arlotta et al 2005), and TLE4 for layer VI (Yao et al 1998). Neurons immunoreactive for each of the three neuronal markers CUX1, CTIP2, and TLE4 were detected in the ectopias (Fig. 3C). Therefore, the ectopic cells in the *Eml3* null FNEs are neurons from deep and superficial cortical layers. These observations suggested that the developmental defect that leads to the FNEs in *Eml3* null mice is present when the first radially migrating neuroblasts of the cortical plate reach the PBM.

In addition, analysis of neuronal marker layering determined that cortical lamination outside of the FNEs is normal (Fig.3C), indicating that neurogenesis and neuroblast radial migration are normal outside of the FNEs.

#### Neuroblasts from the first radially migrating cohort over-migrate into the subarachnoid space

To identify the developmental stage at which the first neuronal over-migration events occur we looked at high spatial and temporal resolution for gaps in the PBM and the presence of over-migrating neurons. Our analysis revealed that the earliest occurrence of a gap in the PBM with neuronal over-migration was in an *Eml3* null embryo with 39 somite pairs (TS17, collected at E11.5; Fig. 4A). Notably, the few tubulin beta-3 immunoreactive neuroblasts observed immediately below the PBM at the 39-somite pair stage represent the first cohort of cells to reach the PBM through radial migration.

**Fig. 4.**
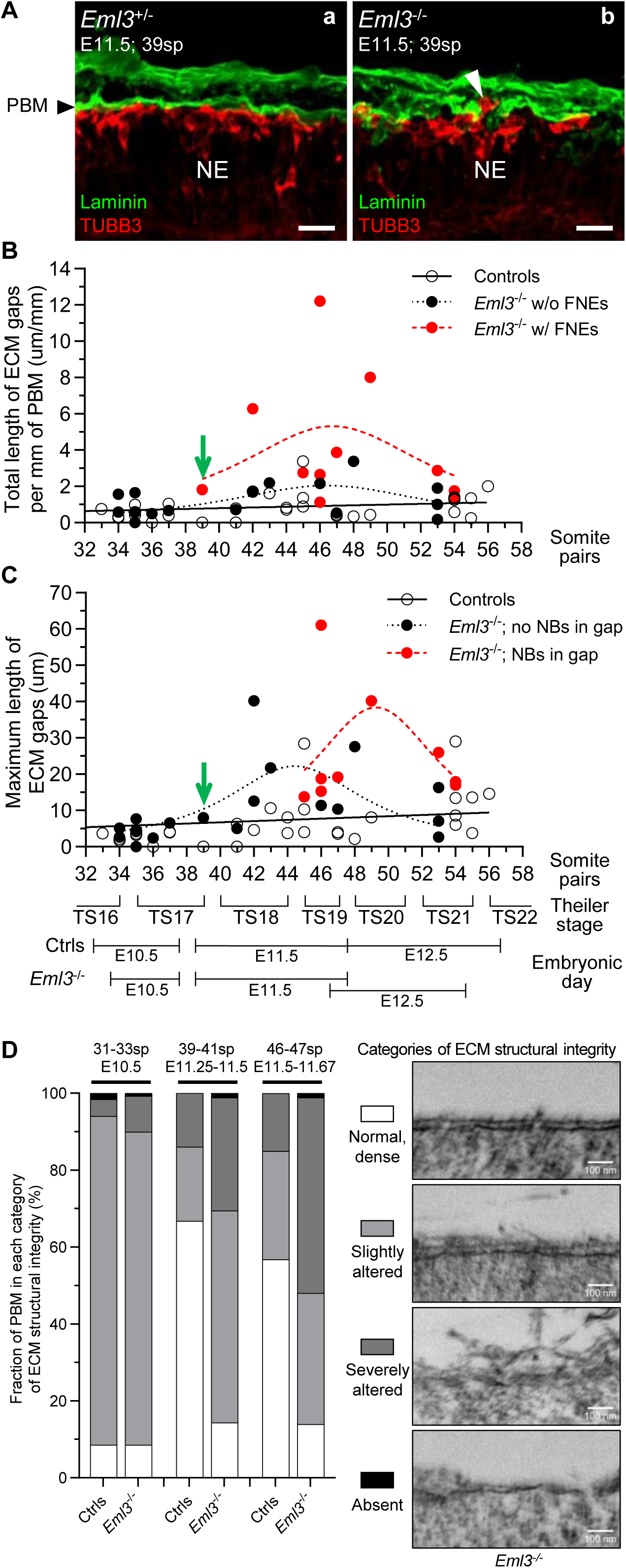
Onset of neuronal over-migration and structural integrity of pial basement membrane ECM. **(A)** Double immunostaining of laminin (green) and tubulin beta-3 (TUBB3, red) at 39 somite pair (sp) embryonic stage (TS17; collected at E11.5). Scale bars, 20µm. **(a)** In a heterozygous brain the PBM is continuous and migrating neuroblasts are observed underneath, whereas in an *Eml3*^-/-^ embryo **(b)** a gap in the ECM and ectopic neuroblasts (arrowhead) were detected. **(B, C)** Coronal head sections of control and *Eml3*^-/-^ embryos at different developmental stages were stained for laminin and tubulin beta-3. The length of gaps in the ECM was measured and the presence of over-migrating neuroblasts (NBs) was noted. The total length of PBM evaluated was measured for normalization. The green arrow points to the *Eml3*^-/-^ embryo that had an FNE at the earliest developmental stage i.e., 39 somite pairs. **(B)** For each embryo, the total length of gaps per mm of PBM analyzed was plotted against the number of somite pairs counted. The *Eml3*^-/-^ embryos with and without FNEs were plotted separately. **(C)** For each embryo, the length of the largest gap observed was plotted against the number of somite pairs counted. For the *Eml3*^-/-^ embryos, the largest gaps with and without over-migrating NBs through the gap were plotted separately. No over-migrating NBs were ever observed in gaps of control embryos. **(D)** Transmission electron microscopy was performed on sections of *Eml3^-/-^* and control embryos (Ctrls; *Eml3^+/+^* and *Eml3^+/-^*) matched by developmental stage. Three stages were analyzed (31-33, 39-41, and 46-47 somite pairs; sp). The embryo ages in days post-coitum are indicated. The total length of PBM was measured in each TEM micrograph. The ECM length was then subdivided into regions with different structural integrity. Normal, dense: sharp, straight, electron-dense ECM. Slightly altered: thicker, diffuse, less dense ECM. Severely altered: delaminating ECM, with electron dense material shedding off a diffuse PBM. Absent: regions with no ECM overlaying the NEP basal end-feet. Representative micrographs shown. At the 31-33sp stage, control ECM was 8.6±7.8% normal (dense), 85.4±8.0% slightly altered, 4.3±3.6% severely altered, and 1.6±1.4% absent, whereas *Eml3^-/-^* ECM was 8.5±7.0% normal (dense), 81.4±2.9% slightly altered, 9.3±8.1% severely altered, and 0.7±1.1% absent. At the 39-41sp stage, control ECM was 66.8±6.3% normal, 19.3±9.0% slightly altered and 13.9±15.2% severely altered, whereas *Eml3^-/-^* ECM was 14.4±6.7% normal, 55.1±23.8% slightly altered, 29.3±30.2% severely altered and 1.2±0.4% absent. At the 46-47sp stage, control ECM was 56.8±9.1% normal, 28.2±4.0% slightly altered and 15.0±5.1% severely altered, whereas *Eml3^-/-^* ECM was 13.9±5.9% normal, 34.1±7.2% slightly altered, 50.8±11.6% severely altered and 1.2±1.6% absent.

#### Increased PBM discontinuity in Eml3 null embryos coincides with the arrival of radially migrating neuroblasts

Over-migration of neuroblasts into the subarachnoid space can occur because of defects in the integrity of the PBM which can no longer halt the migrating neurons (Radmanesh et al 2013, van Reeuwijk et al 2005a). Alternatively, neurons unable to detect or integrate migration stop cues can punch through a normal PBM (Moers et al 2008). The presence of defects in PBM structure prior to the arrival of radially migrating neuroblasts can distinguish between the two possible mechanisms for FNE formation.

As a first step in the characterization of the *Eml3* null PBM, we immunostained for laminin, collagen IV, α-dystroglycan, and integrin α6. No major differences were observed (Supp. Fig. 2).

To further characterize the PBM of *Eml3* null embryos, we quantified gaps in the laminin immunoreactive PBM in embryos at developmental stages between 33 and 56 somite pairs (Fig. 4B and 4C). The cross-sectional length of each gap was measured. The total length of all the gaps observed in the PBM of an embryo (μm per mm of PBM analyzed, a measure of gap burden) was calculated and plotted against the number of somite pairs counted for that embryo (Fig. 4B). The cross-sectional length of the largest gap per embryo (maximum cross-sectional length in μm) was also plotted against the number of somite pairs (Fig. 4C). In control brains, PBM gaps were rare (average gap burden of 1.1 μm per mm of PBM; Fig. 4B) and small (average maximum gap size below 10 μm; Fig. 4C) at all developmental stages examined. The total and maximum size of gaps did not significantly differ between control and *Eml3* null embryos of fewer than 39 somite pairs (Fig. 4B and 4C). However, in embryos with more than 39 somite pairs, the total length of gaps per mm of PBM was significantly larger in *Eml3* null embryos than in control embryos (Fig. 4B). The gap burden in μm per mm PBM peaked at the 47-somite pair stage in *Eml3* null embryos and was larger in *Eml3* null embryos with FNEs (about 5.3 μm gap per mm of PBM) than in *Eml3* null embryos without FNEs (2.0 μm gap per mm of PBM). When maximum gap size was analyzed, *Eml3* null embryos had significantly larger gaps than control embryos after the 39-somite pair stage (Fig. 4C). In *Eml3* null embryos, some of the largest PBM gaps did not have over-migrating neuroblasts. Indeed, in the *Eml3* null embryo with the earliest observed FNE, the over-migrating neuroblasts were not within the largest gap found in that embryo. Best-fit curves for maximum ECM gap size in *Eml3* null embryos peaked at 22 μm at the 44-somite pair stage when there were no ectopic neuroblasts visible in the gap. However, when there were ectopic neuroblasts within the gap, it reached 38 μm at the 49-somite pair stage (Fig. 4C).

Thus, *Eml3* null embryos have larger gaps and a higher gap burden than control embryos. The gap burden is higher in *Eml3* null embryos that have FNEs than in *Eml3* null embryos that do not have FNEs. Also, gaps with over-migrating neuroblasts are larger than gaps that do not have over-migrating neuroblasts. Importantly, PBM gaps without over-migrating neuroblasts are larger in *Eml3* null than in control embryos. Hence, the *Eml3* null PBM is defective in the absence of over-migrating neuroblasts and the hypothesis in which neuroblasts punch through an intact PBM in *Eml3* null developing cortex can therefore be excluded.

#### Structural abnormalities in the ECM of the Eml3 null pial basement membrane

Since the differences in gap size and gap burden are not yet significant when the first FNEs appear, we hypothesized that neuroblasts pass through the PBM when the structural defects are not detectable through laminin immunostaining. We therefore looked for subtle structural abnormalities of the PBM before the onset of FNEs. To analyze the structure of the PBM at a higher resolution, transmission electron microscopy (TEM) was performed on sections of control and *Eml3* null embryos. Brains were analyzed at three developmental stages: at 31-33 somite pairs, before the onset of FNEs; at 39-41 somite pairs, the onset of FNEs; and at 46-47 somite pairs, the stage when the gap burden was largest in *Eml3* null embryos. The total length of PBM was measured in each TEM micrograph and subdivided into regions with different structural integrity (Fig. 4D). A sharp, straight, electron-dense ECM was categorized as normal and dense. A thicker, diffuse, less dense ECM was categorized as slightly altered. A delaminating ECM, with electron-dense material shedding off a diffuse PBM was categorized as severely altered. Regions with no ECM overlaying the neuroepithelial basal endfeet was categorized as absent. At the 31-33 somite pair stage, the immature PBM ECM is not yet fully condensed, therefore for both genotypes most of the ECM was categorized as slightly altered given its low density. Percentages of the PBM that were assigned to each category of structural integrity were significantly different between *Eml3* null and control mice at two of the developmental stages studied. Although no differences in the ECM morphology were observed at the 31-33 somite pair stages, a higher proportion of structurally disrupted ECM in *Eml3* null embryos both at the 39-41 somite pair and at the 46-47 somite pair stages was observed (Fig. 4D). Importantly, the morphology and spacing of the basal endfeet of neural progenitors were not different in *Eml3* null and control embryos according to the TEM micrographs at the stages studied. Thus, we conclude that, at the onset of FNEs, the structural integrity of the PBM is compromised, with a structurally abnormal ECM.

### EML3 is expressed in the tissues that form and maintain the PBM

To determine if the EML3 protein is expressed in the cells that secrete and stabilize the ECM of the PBM, EML3 immunostaining was performed on E10.5 embryo heads (Fig. 5A). E10.5 embryos with 35-37 somite pairs are at the developmental stage that immediately precedes the onset of PBM defects and FNEs. The antibody specificity was validated with an *Eml3* null littermate. At this stage, the EML3-specific signal is ubiquitous, including the PBM-forming mesenchyme and neuroepithelial cells. Neuroepithelial cells span the entire cerebral cortex at this stage, however the EML3 signal is strongest near the ventricles, suggesting a preference for an apical subcellular localization in those polarized cells.

**Fig. 5.**
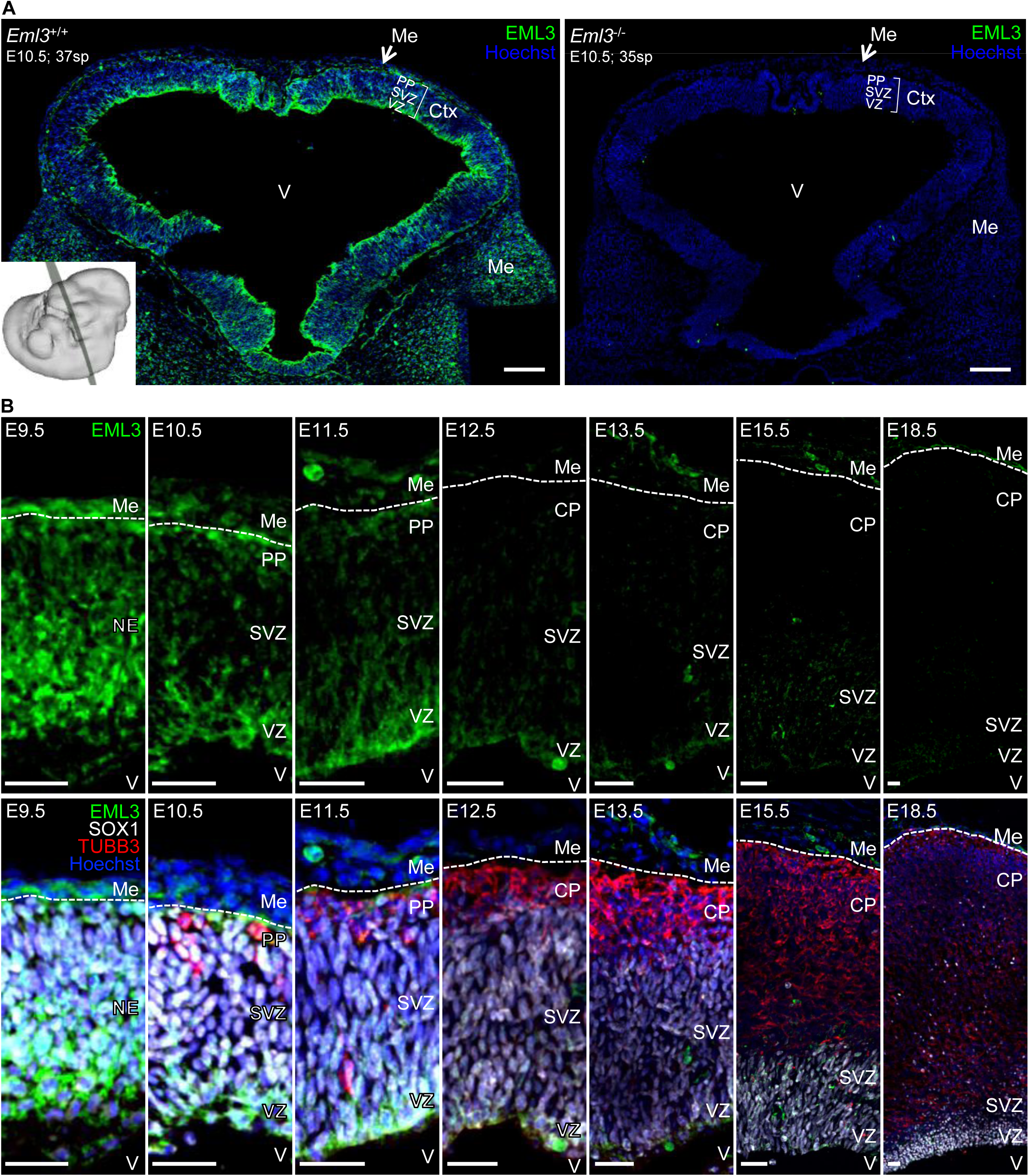
EML3 is expressed in the tissues that form and maintain the PBM. **(A)** EML3 protein expression (green) as determined by indirect immunofluorescence on a coronal cryosection of an E10.5 *Eml3*^+/+^ embryo head. This developmental timepoint, 37 somite pairs, immediately precedes appearance of the first FNE in *Eml3*^-/-^ embryos (at 39 somite pairs). An *Eml3*^-/-^ littermate with 35 somite pairs was included as control for antibody staining specificity. Ctx, cerebral cortex; Me, mesenchyme; PP, preplate; SVZ, subventricular zone; VZ, ventricular zone; V, ventricle. Hoechst nuclear stain (blue). Scale bars, 100µm. **(B)** EML3 protein expression (green) during forebrain development as determined by indirect immunofluorescence on E9.5-E18.5 *Eml3*^+/+^ embryo cryosections. In the bottom panels, the same EML3-stained section is shown with additional immunostaining for markers of neural progenitor cells (SOX1, white) and of post-mitotic neuronal cells (TUBB3, red), as well as Hoechst nuclear stain (blue). The dotted line indicates the location of the PBM which separates mesenchyme (Me) and cerebral cortex whose cell composition changes during development. At E9.5 the cortex is spanned by neuroepithelial cells (NE). At E10.5 and E11.5, the neuroepithelial progenitors are differentiating into radial glial cells and the cortex can now be divided into ventricular zone (VZ), subventricular zone (SVZ), and the post-mitotic neurons of the preplate (PP). At E12.5-E18.5, radially migrating neurons are populating the region below the PBM and form the developing cortical plate (CP). V, ventricle. Scale bars, 25µm.

To further localize the EML3 protein in the PBM-forming cells during forebrain development, coronal sections of embryonic mouse heads at E9.5-E18.5 were stained with anti-EML3 antibody. EML3 immunoreactivity was detected in the developing brain cortex and mesenchyme at E9.5-E18.5 (Fig. 5B) but the levels were highest at the stages that precede the onset of FNEs (E9.5 and E10.5). EML3 expression levels decrease over time, with a marked decrease between E11.5 and E12.5, when neuroepithelial cells are completing differentiation into radial glia (Gotz & Huttner 2005, Noctor et al 2008). By E18.5, EML3 protein expression was limited to the thin mesenchymal cell layer and the ventricular zone (VZ) of the brain cortex, where SOX1-positive precursors are still present. In both neuroepithelial (E9.5-E11.5) and radial glial cells (E11.5-E18.5), the subcellular localization of the EML3 protein was predominantly apical. In neuroepithelial cells (E9.5-E11.5), the second strongest EML3 signal was in the basal endfeet, which are part of the PBM. At E10.5 and E11.5, co-staining with a marker for post-mitotic neurons, TUBB3, revealed that EML3 protein is also expressed in early migrating neurons, albeit at lower levels than in the neural precursors (Fig. 5B). EML3 is therefore expressed in the cells that secrete and stabilize the ECM components of the PBM and the timing of the EML3 expression overlaps with the onset of the *Eml3* null PBM defects.

### EML3 protein interactions

To establish the molecular mechanism by which the absence of EML3 protein leads to focal neuronal ectopias and intrauterine growth restriction of mouse embryos, we searched for proteins that interact with EML3. Taking advantage of *Eml3* null mice to control for non-specific interactions, we used co-immunoprecipitation coupled to mass spectrometry (coIP-MS) to find EML3-interacting proteins *in vivo* in tissues that could be relevant to the phenotypes that we observed. Immunoprecipitations with anti-EML3 antibody were performed on E15.5 *Eml3^+/+^* and *Eml3^-/-^* whole embryo lysates (three biological replicates per genotype) and on E12.5 *Eml3^+/+^*and *Eml3^-/-^* head lysates (two biological replicates per genotype). Additionally, as defects in extra-embryonic tissues are often associated with intrauterine growth restriction, immunoprecipitations were performed on E12.5 placenta lysates (two biological replicates). Table 3 displays the proteins that are immunoprecipitated with the EML3 antibody from each of the tissues of interest. As expected, EML3 is the most abundant protein identified in the immunoprecipitates. The most abundant co-immunoprecipitate of EML3 in E12.5 head samples is the neuron-specific tubulin beta-3 (TUBB3). Five 14-3-3 proteins were identified as co-immunoprecipitating with EML3. When all 14-3-3 spectral counts from the five paralogs are combined, the 14-3-3 protein family is the most abundant EML3 co-immunoprecipitate. Among the 14-3-3 proteins, 14-3-3 theta was the most abundant EML3 co-immunoprecipitate. EML3 contains the ideal binding site for the cytoplasmic dynein light chain (DYNLL) proteins (Rapali et al 2011) and we identified DYNLL1 as one of the most abundant EML3 co-immunoprecipitates. The association of the EML3 protein with DYNLL1, 14-3-3 epsilon (YWHAE), and 14-3-3 gamma (YWHAG) was verified through co-immunoprecipitation in transfected cells (Fig. 6 and data not shown).

**Fig. 6.**
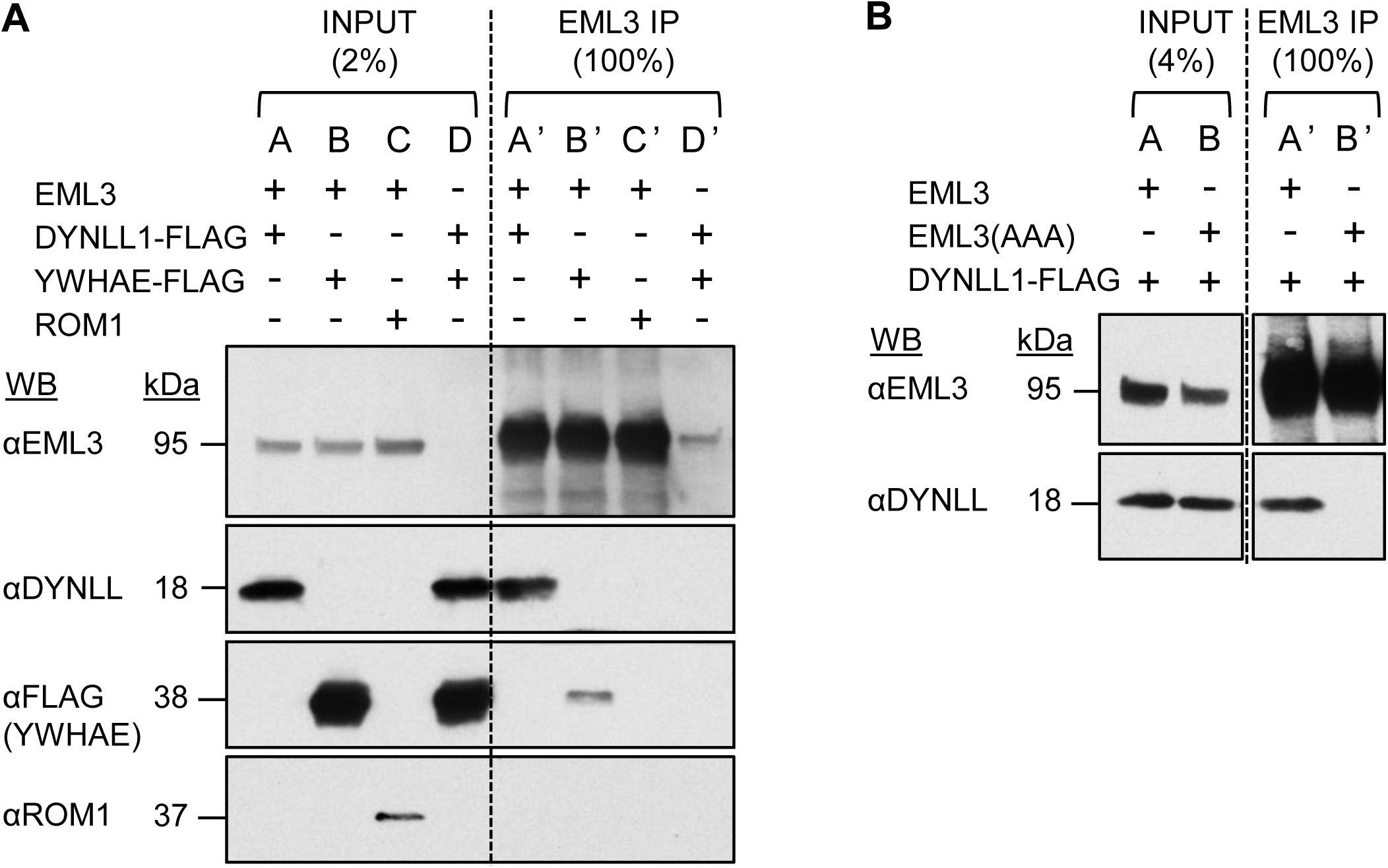
Verification of EML3 protein interactions in co-transfected cells. HEK293T cells were transfected with plasmids encoding for the indicated full-length proteins. A fraction of the cell lysates was immunoblotted for verification of protein expression (INPUT lanes, 2% or 4% of the total lysate) and the remainder was used for immunoprecipitations with EML3 rabbit polyclonal antibody (EML3 IP lanes). HEK293T cells express low amounts of endogenous EML3 protein. **(A)** EML3 interacts with DYNLL1 and YWHAE proteins in co-transfection experiments. DYNLL1 co-immunoprecipitates with co-expressed EML3 protein in lane A’. YWHAE co-immunoprecipitates with co-expressed EML3 protein in lane B’. ROM1 does not co-immunoprecipitate with co-expressed EML3 protein in lane C’. DYNLL1 and YWHAE are not immunoprecipitated by the EML3 antibody when no EML3-expressing construct is co-transfected into the cells in lane D’. **(B)** Mutating the TQT86 motif of the EML3 protein to AAA abolishes binding to DYNLL1. DYNLL1 co-immunoprecipitates with co-expressed EML3 protein in lane A’. DYNLL1 does not co-immunoprecipitate with co-expressed EML3^TQT86AAA^ mutant protein in lane B’.

**Table 3.**
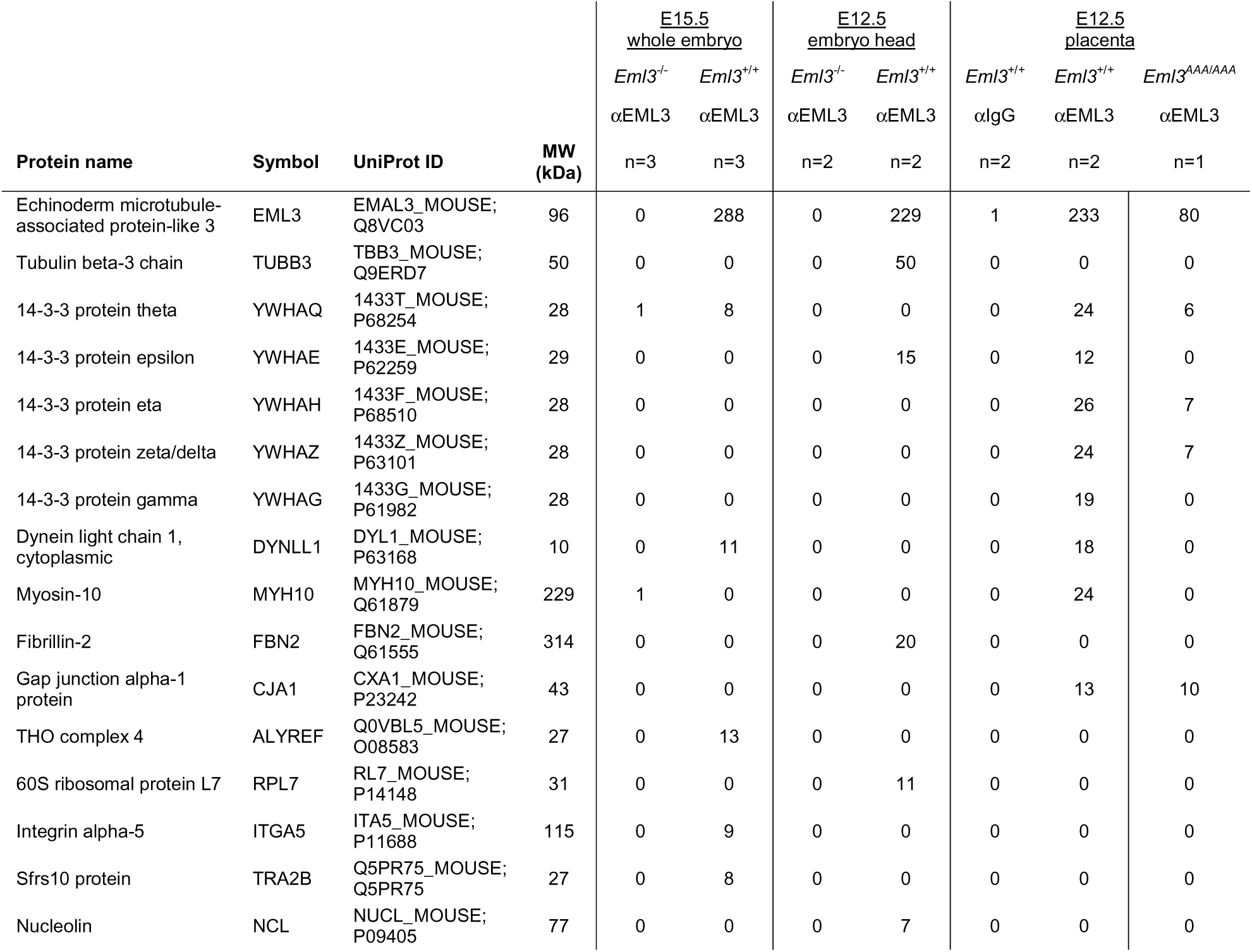
EML3 protein coIP-MS data from three different tissues.

To define what biological processes are dependent on an EML3-DYNLL interaction, we generated mice in which the EML3 DYNLL-binding motif was mutated to abolish binding. In EML3, the DYNLL-binding site is a TQT motif at position 86 of the protein sequence (Rapali et al 2011). An expression construct in which the EML3 TQT motif was replaced by alanines (TQT86AAA) was generated. Figure 6B shows a co-immunoprecipitation experiment performed in cells transfected with wild type EML3 and the TQT86AAA variant. The TQT86AAA mutation abolished EML3-DYNLL binding. A mouse line carrying the TQT86AAA mutation was generated. DYNLL was not co-immunoprecipitated with EML3 in lysates from homozygous *Eml3*^TQT86AAA^ E12.5 placentas (*Eml3^AAA/AAA^*) in coIP-MS experiments (Table 3). Remarkably, mice homozygous for the *Eml3*^TQT86AAA^ allele developed normally, with no intrauterine growth restriction and no focal neuronal ectopias in the developing cortex. The mutant mice survived past weaning age, appeared normal, were fertile and had a normal lifespan without apparent health issues. Thus, the DYNLL1 binding site and EML3-DYNLL direct interaction is dispensable for EML3 functions identified in *Eml3* null mice.

## DISCUSSION

In this study, we show that loss of EML3 results in focal neuronal ectopias with over-migration of neurons through a defective PBM. EML3 is the second member of the EML protein family, with EML1, whose absence leads to cortical brain malformations. However, the cortical malformation caused by the absence of EML3 contrasts with the subcortical band heterotopias observed in *Eml1* null mice (Bizzotto et al 2017, Kielar et al 2014).

### Intrauterine growth restriction and delayed embryo development in *Eml3* null mice

In addition to the cortical brain defect, *Eml3* null mice have a delay in embryonic development, have intrauterine growth restriction, and die perinatally. The growth restriction and somite count differences observed in *Eml3* null embryos become apparent at E8.5. No size or stage differences were seen at E7.5 nor in pre-implantation embryos. We observe no differences in staining for cell cycle (pHH3 and Cyclin B1) and apoptotic cell death (cleaved caspase 3) markers in E8.5 embryonic and extraembryonic tissues (data not shown). We did not pursue further the elucidation of the molecular mechanism responsible for the growth restriction and development delay of *Eml3* null embryos.

Nevertheless, this delay had to be considered for the characterization of *Eml3* null focal neuronal ectopias. Consequently, neurodevelopmental analyses were performed on developmentally/stage-matched embryos because comparison of *Eml3* null and control littermates displayed differences in cortical development that were a consequence of the developmental delay.

### Role of EML3 in the integrity of the PBM and ECM remodeling

Our electron microscopy analyses demonstrate that the absence of EML3 results in structural defects of the PBM that precede breaching by radially migrating neuroblasts. At the TEM level, we found that the PBM of the dorsal telencephalon of *Eml3* null mice was uniformly disorganized and less dense than that of control mice (Fig. 4D). However, the FNEs, which are the macroscopic structural anomaly, occur focally, are relatively few, and occur in random locations within the dorsal telencephalon (Figs. 2A and 2C). Given that the first FNEs occurred when the first radially migrating neurons reached the already defective PBM, we propose that the abnormal PBM in *Eml3* null embryos is sporadically disrupted by the first migrating neurons of the cortical plate, creating a permanent breach through which subsequent neurons can also migrate into the adjacent subarachnoid space.

Interestingly, basement membranes other than the PBM, such as that of the skin, blood vessels, muscle, or retina were not affected in *Eml3* null embryos (data not shown). This suggests that the absence of EML3 uniquely affects the integrity of the PBM. One possible explanation is that the component or pathway defective in *Eml3* null mice is redundant in basement membranes other than the PBM.

The small number and size of FNEs in *Eml3* null mice is comparable to the GPR56 (Li et al 2008) and Gα_12/13 (_Moers et al 2008_)_ mouse models of COB. GPR56 is a receptor for the ECM component collagen III and couples to the Gα_12/13_ family of G proteins for activation of the RhoA pathway upon ligand binding (Luo et al 2011). However, since many different receptor-ligand pairs exist, there is redundancy in the binding of ECM components to radial glial endfeet. In fact, GPR56 appears to play a lesser role in PBM formation and maintenance than other ECM receptors. For example, *Gpr56* null mice have less severe COB phenotypes than mice with mutations in dystroglycan (Geis et al 2013, Godfrey et al 2007, Myshrall et al 2012, Riemersma et al 2015). Thus, as seen for GPR56, the molecular or cellular function of the EML3 protein in PBM formation or maintenance may be partly redundant. An additional level of redundancy for EML3 function has been unveiled by transferring *Eml3* null alleles into the CD-1 outbred mouse strain. The resulting CD-1 congenic *Eml3* null mice are indistinguishable from control littermates, with no perinatal lethality, intrauterine growth restriction, nor FNEs (data not shown). Thus, CD-1 outbred mice have modifier genes that compensate for the absence of EML3, whereas inbred strains of mice 129X1 (data not shown) and C57BL/6 do not.

### Identifying the tissue(s) in which EML3 is essential for cortical brain formation

The PBM is made up of an ECM and the basal endfeet of radial glial processes. Meningeal fibroblasts in the mesenchyme contribute to the PBM by secreting and organizing most of the basement membrane constituents (Sievers et al 1994). Cell surface receptors, such as integrins and dystroglycan, on basal endfeet of neuroepithelial cells and radial glia, orchestrate the assembly of the ECM (Schwarzbauer 1999). Thus, the integrity of the PBM relies on both the meningeal fibroblasts and the neuroepithelial or radial glial cells of the cerebral cortex. In this study, we establish the role of EML3 in regulating PBM integrity by demonstrating that 1) loss of EML3 in mice results in defective PBM and focal neuronal ectopias, a cobblestone-like cortex, and 2) EML3 is expressed in both tissues that contribute to PBM formation and maintenance, namely the meningeal mesenchyme and the neuroepithelium, at the time when the PBM is forming (Fig. 5). However, the tissues in which EML3 expression is essential for normal PBM formation and maintenance remain to be determined. With the conditional potential of the *Eml3^fl/fl^* mice that we have generated (Supp. Fig. 1), it will be possible to delete EML3 in specific tissues and at specific developmental stages.

### Differences in the spatio-temporal expression of EML3 and EML1 proteins

Our study indicates that EML1 and EML3 proteins affect neuroblast migration in opposite directions, with EML1 deficiency resulting in subcortical band heterotopia with under-migration of neuroblasts and EML3 deficiency resulting in cobblestone brain malformation with over-migration of neuroblasts. Importantly, absence of neither EML1 nor EML3 results in a neuroblast-intrinsic cell motility defect. In *Eml1* null brains, radial glia have been defined as the defective tissue whereas in *Eml3* null brains, the two main candidate tissues are neuroepithelium and mesenchyme. Whereas the EML3 protein is present in both of the PBM forming tissues, *Eml1* mRNA was shown to be expressed in neural progenitors and not in meningeal mesenchyme (Kielar et al 2014). The timing of the expression of EML1 and EML3 also differs, with EML3 being expressed at its highest levels until E11.5, while *Eml1* mRNA expression was observed in the radial glia, intermediate progenitors, neuroblasts and mature neurons starting at E11.5. Tissue and developmental stage specificity in the expression of the two EML proteins may account at least in part for the contrasting effects of their absence on neuroblast radial migration.

### Analysis of microtubule-dependent cellular processes in the developing *Eml3* null brain

EML3 is a microtubule-binding protein with no documented link to PBM formation or maintenance. Identifying the cellular and molecular mechanisms by which the absence of EML3 protein results in a defective PBM may therefore unveil a novel pathological mechanism for COB.

The EML3 protein was first isolated from HeLa cell microtubule preparations (Tegha-Dunghu et al 2008) and it was then shown to colocalize with microtubules in cell cultures (Luo et al 2019, Tegha-Dunghu et al 2008). Indeed, our pulldown experiments from embryo lysates supported their interaction in vivo. Moreover, we have determined that EML3 binds polymerized microtubules directly *in vitro* (TIRF microscopy, data not shown). Microtubule-associated proteins (MAPs) play a crucial role in regulating the dynamics and organization of microtubules in cells (Alfaro-Aco & Petry 2015, Bodakuntla et al 2019). The absence of EML3 protein may therefore alter microtubule dynamics or organization. However, staining of *Eml3* null and control cortices with the microtubule markers alpha-tubulin, acetylated alpha-tubulin, and tubulin beta 3 at all embryonic ages studied revealed no differences in the localization or organization of microtubules (data not shown).

Microtubules play important roles in cellular processes such as cell division, cell polarity, cell motility, and trafficking of organelles (Goodson & Jonasson 2018, Logan & Menko 2019). Microtubule-dependent processes were therefore considered as possible cellular mechanisms by which the absence of EML3 leads to COB. No defects in cell division were observed in the developing *Eml3* null brains. In particular, no significant differences were observed between *Eml3* null and control littermates in the mitotic index nor in the percentage of Cyclin B1 positive cells in the neuroepithelium and in the head mesenchyme in E8.5-E11.5 embryos (data not shown). Also, no differences in the angle of mitotic spindles in apical progenitors of stage-matched *Eml3* null and control embryos were observed (data not shown). We ruled out that the *Eml3* null PBM defects are a consequence of a loss of neural precursor polarity by staining neuroepithelial and radial glial cells in E9.5-11.5 *Eml3* null and control brain sections with apical and basal markers (data not shown). In areas of the *Eml3* null developing telencephalon with an intact basement membrane, neurons migrated to their appropriate destination and the cortex is layered normally (Fig. 3C). The motility of radially migrating neuroblasts therefore appears to be normal in the absence of PBM breaches. Additionally, we saw no differences in the staining pattern for ECM components between *Eml3* null and control brains (data not shown) suggesting no major deficiencies in the secretion of main PBM ECM components. Our observations are therefore not consistent with a direct role of the EML3 protein in microtubule dynamics and organization. An alternative function for MAPs, besides a direct role in microtubule dynamics and organization, is to recruit other proteins to microtubules so they can perform their biological functions.

### EML3 protein interactions in relevant tissues and significance for FNE formation

Published high-throughput experiments using cultured cell lines have yielded many potential EML3 interacting proteins, among which the following have been observed in more than one experiment: dynein light chain proteins DYNLL1 and DYNLL2 (Hutchins et al 2010, Huttlin et al 2021); 14-3-3 proteins YWHAB, YWHAG, YWHAH, YWHAQ, and YWHAZ (Ewing et al 2007, Huttlin et al 2021); NEK kinases NEK1, NEK6, NEK7 and NEK9 (Buljan et al 2020, Ewing et al 2007, Golkowski et al 2023, Huttlin et al 2021); and beta-tubulins TUBB2A, TUBB2B, TUBB4B, TUBB8 (Huttlin et al 2021).

As determined previously for EML1 and EML2 (Richards et al 2014), we confirmed that EML3 binds soluble tubulin proteins. We also confirmed the interaction of EML3 with DYNLL and 14-3-3 proteins and determined with co-transfection assays that the pairwise interactions with EML3 are direct (Table 3, Fig. 6, and data not shown). Moreover, using TIRF microscopy with purified proteins, we determined that EML3 can recruit DYNLL to microtubules (data not shown). Interestingly, we found that the EML3-DYNLL interaction was dispensable for the function of EML3 since the mouse line carrying the mutation in the interacting domain had no impact on mouse development.

YWHAE is the only 14-3-3 protein that we identified as an EML3 interactor in the developing brain and *YWHAE* null mice have phenotypes similar to those of *Eml3* null mice, such as intrauterine growth restriction, perinatal death, and cortical brain malformations (Toyo-oka et al 2003). *YWHAE* null embryos were found to have a thin cortex and slow migration of neuroblasts without inversion of cortical layers.

Luo *et al*. determined that EML3 regulates mitotic spindle assembly by recruiting the augmin protein complex and the gamma-tubulin ring complex to spindle microtubules (Luo et al 2019). EML3 was shown to interact, through its C-terminal domain, with several subunits of the augmin complex as well as with gamma-tubulin. We did not co-IP any of the subunits of the augmin complex, nor gamma-tubulin, in our coIP-MS experiments. However, these interactions occur specifically during mitosis. Since we did not enrich tissue lysates for mitotic cells in our coIP-MS experiments, and mitotic cells represent ∼5% of cells in these tissues, these interactions might have been below the threshold for detection. Another documented EML protein interaction that is transient and cell cycle dependent is with NEK kinases (Adib et al 2019) and we did not identify the NEK kinases as interactors in the embryo tissues.

### Clinical relevance

Mutations in *EML1* have been linked to band heterotopia in humans (Kielar et al 2014, Shaheen et al 2017). To date, no *EML3* mutations of clinical relevance have been identified. Given that *Eml3* null mice are a model of cobblestone brain malformation and that approximately one-third of non-syndromic COB cases have no known etiology, we analyzed a panel of DNA samples from 38 patients with COB and no known causative mutations in any of the known dystroglycanopathy genes. No *EML3* coding mutations were seen in those samples. Since COB and polymicrogyria (PMG) have phenotypic and etiological similarities, we also analyzed a panel of DNA samples from 15 PMG patients and found no *EML3* coding mutations. Defining the clinical relevance for the *EML3* gene will require further testing for non-coding regulatory mutations as well as testing additional COB samples. The phenotypic spectrum for genetic analyses could also be expanded; given the phenotypes observed in our *Eml3* null mice the *EML3* gene may have clinical implications for intrauterine growth restriction (IUGR).

### Conclusion

The microtubule binding protein EML3 is required for mammalian embryonic growth and cortical brain development. *Eml3* null mice exhibit intrauterine growth restriction with delayed embryo development and die perinatally. In addition, *Eml3* null mice have focal neuronal ectopias, due to neuroblasts migrating past a defective PBM into the sub-arachnoid space, phenocopying the cobblestone brain malformation condition. We demonstrated that the structural integrity of the PBM is affected by the time that the first radially migrating neuroblasts reach the marginal zone of the cortical plate.

Although the EML3 protein was reported to be required for mitotic progression in cancer cell lines, we found no differences in mitotic and other cell cycle markers in *Eml3* null embryo tissues. Further cellular and molecular investigations will be necessary to deepen our understanding of how the EML3 protein controls, either directly or indirectly, neuroblast radial migration.

## MATERIALS & METHODS

### Generation of Eml3 null mice

*Eml3* null mice (*Eml3*^tm1e(EUCOMM)Wtsi^) were generated from ES cell line clone EPD0751_4_D11 made by the European Conditional Mouse Mutagenesis Program (EUCOMM) and purchased from the European Mouse Mutant Cell Repository. See Skarnes et al., 2011 for the general design of the lines. Specifically, the ‘knockout-first’ allele of *Eml3* was constructed by inserting an IRES:lacZ trapping cassette and a floxed promoter-driven neo cassette into *Eml3* intron 10 based on NCBI Reference Sequence gene NM_144872. We obtained JM8A3.N1 ES cells at passage 5 (C57BL/6N mouse strain, male genotype) that were heterozygous for the targeted mutation. The ES clone contains a KO-first reporter-tagged insertion with conditional potential (allele *Eml3*^tm1a(EUCOMM)Wtsi^). The nucleotide sequence of that allele is publicly available as GenBank JN963812.

ES cells were injected into C57BL/6N blastocysts at the Goodman Cancer Center Transgenic Core Facility and one chimeric male with germline transmission of the targeted *Eml3* allele was imported into our facility. However, our mice did not carry the distal loxP site and are thus devoid of conditional potential and therefore represent an *Eml3*^tm1e(EUCOMM)Wtsi^ allele according to EUCOMM nomenclature (Supp. Fig, 1B). We determined, through PCR analysis and sequencing, that the ES cell cultures contained a minor fraction (about 15%) of cells with the added loxP site within *Eml3* intron 16 in a background of cells with an intact wild type (WT) sequence. Our founder male was therefore derived from the more prevalent cell type in the mixed ES cell population. The primers used to detect the presence of the distal loxP site were: RM173F 5’-GCAGGGAAGTCTAGGAAGGG and RM165R 5’-GGGCTGCTACCAGACTGAAG and yield amplification products of 170 bp in WT alleles or 222 bp when the distal loxP site is present. Mice were backcrossed to C57BL/6J background (Jax: 000664) for at least two generations prior to conducting experiments and have currently been backcrossed for eight generations. The C57BL/6N-derived retinal degeneration *Crb1^rd8^* spontaneous mutation was eliminated from our mice by selecting *Crb1^wt/wt^* progeny at the N2 generation. Experimental mice were produced by intercrossing heterozygotes.

We then proceeded to generate conditional *Eml3* null mice (*Eml3^fl/fl^*) in a step-wise process starting from the constitutive *Eml3* null (*Eml3*^tm1e(EUCOMM)Wtsi^) mice described above. The first step was to introduce a distal loxP site for Cre-directed removal of coding exons. We opted for a CRISPR-Cas9 mediated homology-directed repair (HDR) technique to introduce a loxP site into *Eml3* intron 19 based on NCBI Reference Sequence gene NM_144872 (genomic position 8,940,293 bp according to GRCm38/mm10 assembly). gRNA 5’-ACTGTTAGTGCTTTATCCAGAGG was used in conjunction with the asymmetric reverse strand repair template (single-stranded oligo deoxynucleotides; ssODN) 5’-TGGGGGCCACCCTGGTCTACAAACGCAGGTGACAGAGGGAACCCTGTCTCAGAAACAGAGAATGTAGCCTTAGACATGCCTAGAGcctctgataacttcgtataatgtatgctatacgaagttatgataaagcactaacagtTTCTCATCAGCTGCCCCTC. The CRISPR-Cas9 reagents: 50ng/ul cas9 mRNA (Sigma-Aldrich, CAS9MRNA-1EA), 20ng/ul *in vitro* transcribed gRNA and 100ng/ul ssODN, were microinjected into the cytoplasm of 1-cell stage embryos obtained from *in vitro* fertilization of C57BL/6N females with sperm from heterozygous *Eml3*^tm1e(EUCOMM)Wtsi^ males. One chimeric male was obtained with in-cis insertion of the distal loxP site into the *Eml3*^tm1e(EUCOMM)Wtsi^ allele. Presence of the intron 19 loxP site was detected by PCR amplification with primers flanking the insertion site and yielding an amplification product of 271 bp in the WT allele or 305 bp when the distal loxP site is present: RM226F 5’-CCGGCTGCACCAAGAAG and RM227R 5’-TTCTCACTGACTTGGCTTGG. The PCR products were sequenced for confirmation that the inserted loxP site was intact. *In-cis* insertion of the distal loxP site with the *Eml3*^tm1e(EUCOMM)Wtsi^ allele was verified by *in vitro* recombination with Cre enzyme (NEB Cat#M0298M) followed by diagnostic PCRs with primers flanking the proximal loxP site within *Eml3* intron 10 (RM177F 5’-AAAACCTCCCACACCTCCC) and distal loxP site within *Eml3* intron 19 (RM227R 5’-TTCTCACTGACTTGGCTTGG), yielding an amplification product of 414 bp only when the distal loxP site was inserted *in-cis* with the proximal loxP site and only in the presence of Cre recombinase. Mice were backcrossed to C57BL/6J background (Jax: 000664) for at least two generations and the C57BL/6N-derived retinal degeneration *Crb1^rd8^*spontaneous mutation was eliminated from our mice by selecting *Crb1^wt/wt^* progeny at the N2 generation. The resulting mouse, *Eml3^em1Mci^*, is a gene trap-floxed allele “*Eml3^gt-fl^*” (Supp.Fig. 1C) and is phenotypically indistinguishable from *Eml3^tm1e(EUCOMM)Wtsi^* mice. The second step was to excise the Gene Trapping cassette and thus produce a WT *Eml3* allele with conditional potential (floxed exons 11-19; *Eml3^em1.1Mci^*) that is a floxed allele “*Eml3^fl^*” (Supp. Fig. 1D). Trap-floxed allele *Eml3^gt-fl^* mice were crossed to the Flp1 line (B6.129S4-*Gt(ROSA)^26Sortm1(FLP1)Dym^*/RainJ; Jax: 009086) to remove the gene trap, lacZ reporter, and neomycin selection cassettes between the two FRT sites within intron 10 of the *Eml3* gene. The FLP recombinase gene was then eliminated in the following breeding history.

Exposure to cre recombinase removes exons 11-19 leading to a frameshift and premature stop codon, which is predicted to lead to nonsense mediated decay of the mRNA. Absence of the EML3 protein was confirmed by immunoblotting and indirect immunofluorescence. Crossing *Eml3^fl/fl^* mice with hemizygous B6.FVB*-Tg(EIIa-cre)^C5379Lmgd^*/J mice (JAX: 003724) resulted in germline recombination of the loxP sites and therefore, after removal of the *E2a-Cre* transgene in the following breeding history, an *Eml3* null allele that does not rely on the activity of a gene trap cassette. This new global knockout mouse is named *Eml3^em1.2Mci^* (Supp. Fig. 1E). Mice were backcrossed to C57BL/6J background (Jax: 000664) for at least two generations prior to conducting experiments and have currently been backcrossed for six generations. Experimental mice were produced by intercrossing heterozygotes. These mice have been determined to be phenotypically undistinguishable from *Eml3*^tm1e(EUCOMM)Wtsi^ and from *Eml3^em1Mci^* mice.

We generated a number of tissue-specific *Eml3*-deficient mice by crossing our *Eml3^fl/fl^* mice with different mouse lines expressing the cre recombinase under the control of promoters that are active in cells and at developmental stages with potential relevance to the phenotypes observed in the global KO. Sox1, Mesp1, and Pdgfra cre drivers were used to delete *Eml3* in tissues involved in development of the pial basement membrane. F2 crosses of *Eml3^fl/fl^* mice with hemizygous B6.B6CB-Sox1tm1(cre)Take mice (Sox1-Cre; Riken BRC no. RBRC05065) resulted in *Eml3* deletion specifically in neuroepithelium. ICR.Cg-Mesp1tm2(cre)Ysa/YsaRbrc mice (Mesp1-Cre; Riken BRC no. RBRC01145) were first backcrossed a minimum of five times into the B6J genetic background before being used to delete *Eml3* specifically in mesoderm-derived mesenchyme. B6.Cg-E2fltg(Wnt1-Cre)2Sor/J mice (Wnt1-Cre; JAX: 022501) and B6-Tg(Pdgfra-cre)1Clc/J mice (Pdgfra-Cre; JAX: 013148) were used to delete *Eml3* specifically from neural crest-derived mesenchyme. All three tissue-specific nulls were indistinguishable from *Eml3^wt^* controls. *Sox1*, *Mesp1*, and *Pdgfra* cre driver pairs and the combination of all three were also indistinguishable from *Eml3^wt^* controls. We also used B6.FVB-Tg(Ada*-cre)5Xiay/J cre driver mice (JAX: 036543) that resulted in *Eml3* deletion in placenta. In all cases, despite confirmed tissue-specific absence of the EML3 protein, the resulting mice were phenotypically indistinguishable from *Eml3^wt^* controls.

### Genotyping of Eml3 null mice

Two PCR genotyping strategies were used in parallel for unequivocal genotype determination.

A four-primer PCR strategy yields a 286bp PCR product for the wild-type *Eml3* allele (*Eml3^wt^*), a 792bp PCR product for both the gene trap allele (*Eml3^gt^*) and the gene trap-floxed allele (*Eml3^gt-fl^*), a 550bp PCR product for the floxed allele (*Eml3^fl^*) and a 380bp PCR product for the tissue-specific (*Eml3^fl^;Cre*) and the global (*Eml3^em1.2Mci^*) *Eml3-*null alleles. The four-primer *Eml3* genotyping was done using a touchdown PCR technique with the common reverse primer RM170 5’-ACACCAGGGTGGTCTCAAAC within *Eml3* intron 10, distal the insertion site for the targeting cassette, and the forward primers RM070 5’-TGATGTTCTAGGCAGGATGTTC within *Eml3* intron 10, proximal to the insertion site for the targeting cassette or RM169 5’-CGCCTTCTATGAAAGGTTGG within the inserted Neo cassette, for the wild type (286bp) or mutant (792bp) allele amplifications respectively. The RM070-RM170 primer pair yields a 550bp product for the floxed allele (*Eml3^fl^*). The additional reverse primer RM227 5’-TTCTCACTGACTTGGCTTGG within *Eml3* intron 19, distal of the inserted loxP site, yields a 380bp PCR product in combination with the forward primer RM070 for exon-deleted tissue-specific (*Eml3^fl^;Cre*) and global (*Eml3^em1.2Mci^*) *Eml3-*null alleles.

A three-primer PCR strategy yields a 521bp PCR product for the *w*ild-type *Eml3* allele (*Eml3^wt^*), a 446bp PCR product for both the gene trap allele (*Eml3^gt^*) and for the gene trap-floxed allele (*Eml3^gt-fl^*) and a 786bp PCR product for the floxed allele (*Eml3^fl^*). The tissue-specific (*Eml3^fl^;Cre*) and the global (*Eml3^em1.2Mci^*) *Eml3-*null alleles do not yield a PCR product with this strategy. The three-primer *Eml3* genotyping was done using a touchdown PCR technique with the common forward primer RM070 5’-TGATGTTCTAGGCAGGATGTTC within *Eml3* intron 10, proximal to the insertion site for the targeting cassette, and the reverse primers RM071 5’-CCCACTGGTGACAATACAGC within *Eml3* exon 11 or RM075 5’-GTACCCCAGGCTTCACTGAG within the proximal inserted FRT site and the GeneTrap cassette, for the wild type (521bp) or mutant (446bp) allele amplifications respectively.

### Generation and characterization of Eml3 TQT86AAA mice

*Eml3* TQT86AAA mice (*Eml3*^AAA/AAA^) were generated at The Center for Phenogenomics in Toronto, Canada. A CRISPR-Cas9 mediated homology-directed repair (HDR) technique was used to introduce nucleotide changes c.256_262ACCCAAA>gCCgcAg based on NCBI Reference Sequence gene NM_144872. gRNA 5’- GGTGACCAGAGGCACCCAAA was used in conjunction with the asymmetric reverse strand repair template (single-stranded oligo deoxynucleotides; ssODN) 5’-AGAGCTGGAGGACCATTGCTCAGGCCAGGGGGTCCAGAGGATGGAACAATCTCCAATTCTTCTTCCGcTgcGGcGCCTCTGGTCACCAAGGAAGGGCTGCATGTAGGTGGCAGTCCTGGAGGGACTGTGATGCTGCTGAG. The CRISPR-Cas9 reagents were microinjected into the cytoplasm of 1-cell stage C57BL/6J embryos. Founder chimeric mice carrying the TQT86AAA allele were identified by PCR amplification with primers flanking the targeted site and yielding an amplification product of 542 bp: RM055F 5’- GGGTGCAGGAAGAAGAGATG and RM067R 5’- TGGTGGGACAGAATGAAAAAG. The PCR products were sequenced. *Eml3* TQT86AAA mice were backcrossed to C57BL/6J background (Jax: 000664) for at least two generations. Routine genotyping was carried out with the same PCR amplification followed by *Tau*I restriction digests that yield 477, 36 and 29bp bands in the WT allele and 280, 197, 36 and 29bp bands in the mutant allele. These mice were determined to be phenotypically undistinguishable from *Eml3^wt^* mice.

### Mice

Breeding animals were on a 12-hr light/dark schedule. Food and water were available ad libitum. Timed pregnant females (E7.5 to E18.5) were sacrificed by CO_2_ asphyxiation after isoflurane anesthesia, followed by cervical dislocation. Ages of mice analyzed are given as embryonic day (E), where the presence of a vaginal plug was considered as embryonic day E0.5. Embryos were dissected from the uteri and placed in PBS on ice. Weight of embryos and placentas was measured in an analytical balance after drying off excess fluid. Embryo sizes were measured as area of the section corresponding to the sagittally bisected E8.5-E10.5 embryos or as weights for E11.5-E18.5 embryos. All experiments on adult mice were performed using both male and female littermates and all data shown is pooled from both sexes after verification that there were no sex-dependent differences. Animal care was in accordance with the federal Canadian Council on Animal Care, as practiced by McGill University and the Lady Davis Research Institute.

### Immunoblotting of E15.5 whole embryo lysates

E15.5 whole embryo lysates were prepared under conditions that are compatible with downstream immunoblotting as well as with coIP-MS experiments. Uteri containing E15.5 embryos were dissected out of euthanized pregnant females and placed in a petri dish filled with PBS on ice. Each conceptus was dissected out of the uterus and transferred to a fresh petri with PBS to dissect the embryo out of the amniotic sac. The tip of the tail was collected for genotyping. The embryo was then placed in 1mL of cold 0.5% NP40-containing lysis buffer in a 5ml round bottom tube (20mM Tris-HCl, pH8.0; 200mM NaCl; 1mM EDTA; 0.5% NP-40; protease inhibitors Aprotinin, Leupeptin, Pepstatin A, PMSF; 1mM Na3VO4; phosphatase inhibitors NaF, NaPPi, beta-glycerophosphate). The embryo was then homogenized with two 10-second pulses using a polytron microtip. The homogenate was decanted into a 1.5mL tube and rotated for two hours at 4^0^C. The lysates were then cleared by centrifugation at 21K RCF for 15 minutes at 4^0^C. The supernatant (1.2mL) was aliquoted into fresh tubes and conserved at - 80^0^C. Protein quantification was performed using a microvolume BCA assay normalized against BSA standards.

50ug of each embryo lysate was resolved on denaturing SDS-PAGE gels (4-20% gradient TGX precast gels, BIO-RAD) and transferred onto nitrocellulose membranes (Amersham Protran supported membrane, GE Healthcare) using a Mini-PROTEAN electrophoresis apparatus (BIO-RAD). Membranes were blocked with 5% milk in TBST buffer and incubated with primary antibodies overnight at 4 °C and then HRP-conjugated secondary antibodies for 30 mins at room temperature. The immunoreactivity of proteins was visualized with Amersham ECL Western Blotting Select Detection Reagent (GE Healthcare) using a ChemiDoc Imager (BIO-RAD). GAPDH immunoreactivity was used as loading control.

### Antibodies

Rabbit antibodies to EML3 were raised by injecting rabbits with a KLH-conjugated peptide encoding the C-terminal tail (884-897) of EML3 (Pierce/Thermo Fisher). Crude serum was used at a 1:1,000 (v/v) dilution for immunoblotting. Affinity-purified antibodies were isolated from antisera and used at 1:25 (v/v) in immunofluorescence labelling. Rabbit monoclonal anti-GAPDH was used as a loading control for immunoblots (NEB 5174P; 1:4,000). HRP conjugated secondary antibodies were used for immunoblotting: donkey polyclonal anti-rabbit IgG (Amersham NA934) and sheep polyclonal anti-mouse IgG (Amersham NA931), both used at 1:20,000 (v/v) dilution.

The following primary antibodies were also used for immunofluorescence (dilutions in v/v): rabbit anti-CCNB1 (G2/M-specific cyclin-B1) (NEB/Cell signaling 4138; 1:200), goat anti-collagen IV (Southern Biotech 1340-01; 1:100), rat anti-CTIP2 (chicken ovalbumin upstream promoter transcription factor-interacting protein 2) (Abcam ab18465; 1:500), rabbit anti-CUX1 (Homeobox protein cut-like 1) (Santa Cruz sc-13024; 1:50), mouse anti-glycosylated α-dystroglycan antibody (DSHB U. Iowa IIH6 C4; 1:50), rat anti-integrin α6 (Millipore MAB1378MI; 1:100), rabbit anti-Engelbreth-Holm-Swarm laminin (Millipore-Sigma L9393; 1:500), rabbit anti-PAX6 (Paired box protein Pax-6) (Covance PRB-278P; 1:500), rabbit anti-pHH3 (phospho-Histone H3) (Millipore Sigma H0412; 1:200), mouse anti-pHH3 (phospho-Histone H3) (NEB/Cell signaling 9706; 1:200), goat anti-SOX1 (Transcription factor SOX-1) (Novus AF3369; 1:40), rabbit anti-TBR1 (T-box brain protein 1) (Abcam ab31940; 1:5,000), rabbit anti-TBR2 (Eomesodermin homolog/T-box brain protein 2) (Abcam ab23345; 1:500), rabbit anti-TLE4 (Transducin-like enhancer protein 4) (kind gift from Dr. S. Stifani, McGill University, Montreal; 1:500), and mouse anti-TUBB3 (Tubulin beta-3) (Promega PR-G7121; 1:1,000).

The following secondary antibodies from Invitrogen were used at a 1:500 (v/v) dilution for immunofluorescence: donkey anti-goat AF594 A-11058, donkey anti-mouse AF405 A-48257, donkey anti-mouse AF594 A-21203, donkey anti-mouse AF647 A-31571, donkey anti-rabbit AF488 A-21206, goat anti-rabbit AF660 A-21073, and donkey anti-rat AF488 A-21208.

The following primary antibodies were also used for immunoblotting: rabbit anti-DYNLL (Dynein light chain LC8) (Origene TA303752; 1:2,000), mouse anti-FLAG (Origene TA50011-100; 1:4,000), and mouse anti-ROM1 (Rod outer segment membrane protein 1) (kind gift from Dr. R. Molday, UBC, Vancouver; 1:20).

### Tissue Preparation, Histology, and Immunofluorescence Labelling

Timed embryos were harvested after isoflurane anesthesia, CO_2_ asphyxiation, and cervical dislocation of pregnant dams.

Embryo heads (E9.5-E18.5) were fixed at 4°C using 2% paraformaldehyde-lysine-periodate in PBS and cryoprotected by using graduated sucrose concentrations (10%, 20%, and 30% sucrose-PBS), frozen in OCT (optimal cutting temperature) compound (Tissue Plus, Fisher HealthCare) over dry ice and sectioned on a cryostat. Cryosections (12-16µm) were collected on Superfrost plus slides and were either stained with 0.5% cresyl violet/0.3% acetic acid (Nissl staining) or processed for immunofluorescence labelling, as described below.

Cryosections were blocked in Mouse-on-Mouse blocking solution for at least 1hr. Sections were then incubated in primary antibodies, diluted in Mouse-on-Mouse antibody diluent, for 1hr at room temperature. Following washes with HBS-Triton X-100, primary antibodies were visualized by incubating for 30mins at RT with appropriate fluorophore-conjugated secondary antibodies (Invitrogen; 1:500). For double or triple labeling, antibodies were applied concurrently during primary and secondary antibody steps. Washed sections were counterstained with Hoechst 33258 (Sigma) and mounted with coverslips and Fluoromount-G (Southern Biotech). All images were captured using a Quorum Wave FX SD confocal microscope (Leica) and processed using the Volocity software package or FIJI. Representative photographs were obtained with the same exposure setting for control and *Eml3* null.

For E7.5 and E8.5 embryos, following euthanasia of timed-pregnant female mice, each conceptus was isolated from the uterus and fixed in 4% paraformaldehyde-PBS for 30mins at 4°C. After rinses in PBS, the conceptus was then cryopreserved in 10%, 20%, and 30% sucrose-PBS before embedding in OCT compound. Cryosections were processed as for embryo heads.

For whole brain immunolabelling at E15.5, embryo heads were first degloved and then sectioned horizontally below palate to keep brain intact in cranium. Crania were fixed in DMSO:methanol at - 20°C, then rehydrated for permeabilization in 1% triton X-100, followed by blocking in BSA, and then primary antibody incubation, which was carried out for 5 days at 4°C. Following washes in blocking solution, crania were incubated with secondary antibodies (Invitrogen; 1:200) for 2 days at 4°C. After further washes in blocking solution, crania were dehydrated in graded methanol solutions and finally cleared in benzyl alcohol:benzyl benzoate (BABB) for visualization under a fluorescence stereomicroscope (Zeiss). Smaller regions of the dorsal telencephalon/PBM were then also imaged on a Quorum Wave FX SD confocal microscope (Leica).

### Electron microscopy

Embryo heads obtained from intercrosses of *Eml3* heterozygous mice were dissected and fixed by immersion in 2.5% glutaraldehyde in 0.1M sodium cacodylate buffer, pH 7.4. After fixation for 24-48h, tissues were postfixed in 1% osmium tetroxide in cacodylate buffer, then dehydrated and embedded in Embed 812 resin (EMS). Ultrathin sections were cut and stained with uranyl acetate and lead citrate. The samples were observed and imaged using the transmission electron microscope Tecnai G2 Spirit BioTWIN fitted with a Gatan Ultrascan 4000 camera, at the McGill Facility for Electron Microscopy. For the 31-33sp stage, four biological replicates for each genotype were analyzed. For the 39-41sp and the 46-47sp stages, two biological replicates for each genotype were analyzed at each timepoint.

### coIP-MS in mouse lysates

Immunoprecipitations were performed on E15.5 *Eml3^+/+^*and *Eml3^-/-^* whole embryo lysates (three biological replicates per genotype). Lysates were prepared as described for immunoblotting, using a 0.5% NP40 detergent lysis buffer with protease and phosphatase inhibitors. Immunoprecipitations were also performed on E12.5 *Eml3^+/+^* and *Eml3^-/-^* head lysates (two biological replicates per genotype) and on E12.5 placenta lysates (two biological replicates). E12.5 tissue lysates were prepared with the following changes to accommodate the small tissues. The tissues were processed in a 420uL volume of tissue lysis buffer and were disrupted with a sonicator using five three-second pulses at 40 second intervals. The final tissue lysate volumes were approximately 500uL each.

Co-immunoprecipitations were performed as follows. 3ug of rabbit polyclonal anti-EML3 C-terminal antibody (affinity purified) was bound to 20uL of washed anti-rabbit IgG Dynabeads (Invitrogen) in PBST buffer for one hour at 4°C with rotation. The anti-EML3 bound beads were then added to 1.5mg of tissue lysate for a total constant volume of 750uL adjusted with tissue lysis buffer. The bead-antibody-lysate mixture was rotated overnight at 4°C. The beads were then washed four times for 20 minutes each at 4°C with rotation in a wash buffer (50mM Tris-HCl, pH8.0; 150mM NaCl; 5mM EDTA; 0.1% NP-40; 1:1000 v/v of each of the protease inhibitors Aprotinin, Leupeptin, Pepstatin A, PMSF; 1mM Na3VO4; 1:100 v/v of each of the phosphatase inhibitors NaF, NaPPi, beta-glycerophosphate). The beads were then washed five times for 30 minutes each at 4°C with rotation in fresh 50mM Ammonium Bicarbonate buffer. The beads were finally suspended in a 50ul volume of Ammonium Bicarbonate buffer and frozen at -80°C. The beads were then processed for mass spectrometry at the IRIC proteomics platform. Processing included on-bead Trypsin digestion before loading on mass spectrophotometer.

*Eml3* null tissues served as controls for EML3 coIP specificity. However, since the placenta of *Eml3^-/-^*embryos contains EML3-expressing tissues derived from the *Eml3*^+/-^ mother (data not shown), a control IgG antibody was used on *Eml3*^+/+^ placenta lysates to control for non-specific interactions in those immunoprecipitations. Following mass spectrometry of the digested immunoprecipitates, the spectra obtained for each sample were assigned to peptides and ultimately to proteins. For each protein, the total number of spectra identified in each tissue is shown. For inclusion in the list, a lower cut-off limit of five spectra when all three tissue datasets are combined was used. A minimal enrichment of at least ten-fold in *Eml3*^+/+^ versus either *Eml3*^-/-^ or IgG control samples was used as a criterion for inclusion in the list.

### coIP of proteins overexpressed in cultured cells

The association of the EML3 protein with DYNLL1, 14-3-3 epsilon (YWHAE), and 14-3-3 gamma (YWHAG) was verified through co-immunoprecipitation in transfected cells. HEK293T cell monolayers were transfected with mammalian expression constructs using JetPrime transfection reagent (Polyplus) according to manufacturer instructions. The expression constructs were purchased from Origene (Rockville, MD) and contained mouse cDNA inserts for untagged *Eml3* (MC201737), *Dynll1*-FLAG (MR219424), *Ywhae*-FLAG (MR203269), untagged *Rom1* (MC205489), or the custom made *Eml3^TQT86AAA^*construct (MC201737 MUTANT). The expression vectors used were pCMV6-Kan/Neo for untagged constructs and pCMV6-Entry for Myc-DDK/FLAG tagged constructs. After 48 hours, the cells were washed three times with ice-cold PBS and then lysed by adding 1mL of ice-cold IP lysis buffer (50mM Tris-HCl, pH7.4; 150mM NaCl; 1mM EDTA; 1% NP-40; protease inhibitors Aprotinin, Leupeptin, Pepstatin A, PMSF; 1mM Na3VO4; phosphatase inhibitors NaF, NaPPi, beta-glycerophosphate). The lysate was transferred into 1.5 mL microtubes. After an incubation on ice for 30 minutes, the lysates were cleared by centrifugation at 21K RCF for 15 minutes. The supernatants were then aliquoted into new tubes. The total yield is typically 1.5 mg of protein.in a 1.25 mL volume.

2ug of affinity purified EML3 antibody was added to 300 uL aliquots of the transfected cell lysates and incubated overnight at 4°C with rotation. 80 uL of washed anti-rabbit IgG Dynabeads (Invitrogen) were then added to the immunoprecipitation samples and rotated for three hours at 4°C. The bead complexes were then washed three times in a Ca+2 and Mg+2 -free PBS supplemented with 0.1% BSA and 2mM EDTA, pH7.4. The bead complexes were then resuspended in 12 uL of 2X sample loading buffer and boiled for five minutes. The eluates (immunoprecipitates) were then loaded on denaturing SDS-PAGE gels (4-20% gradient TGX precast gels, BIO-RAD) in parallel with small aliquots from the original cell lysates (input). Immunoblotting was performed as indicated above.

### Statistical analysis

Statistical analysis was performed using GraphPad Prism. Data are shown as mean ± s.d. or as mean of means ± s.e.m. The variance was estimated for each set of data and the significance test was adapted accordingly. All tests were two-sided. Comparisons of means in two groups were made using either an unpaired Student’s t-test or with a Welch’s t-test to correct for unequal variances.

Comparisons of means in three groups were made using non-parametric one-way ANOVA followed by Dunn’s pairwise tests.

## ACKNOWLEDGMENTS

This work was supported by a CIHR-Canada Research Chair and the Jewish General Hospital Foundation. SB has been supported by CIHR project grant PJT-189995 and NSERC discovery grant RGPIN-2024-05603. SS also received CIHR funding.

We would like to thank the following individuals: Marilene Paquet and Jean-Martin Lapointe for pathology expertise, Christian Young and Mathew Duguay from the Imaging Core of the Lady Davis Research institute, Veronique Michaud from the Mouse Phenotyping Core of the Lady Davis Research institute, Jeannie Mui at the McGill Facility for Electron Microscopy Research, Julie Lamarche and Jessica Di Giovanni for their help with mouse colony management. Drs. Michel Cayouette and Loydie Jerome-Majewska for helpful advice and support.

## AUTHOR CONTRIBUTIONS

I.C. and E.D. designed and performed experiments, analyzed and prepared the data for publication, co-wrote the manuscript. Specifically, I.C. setup timed matings and collected embryos, phenotyped embryos and post-natal mice, processed tissues and performed indirect immunofluorescence and electron microscopy. I.C. discovered and characterized the focal neuronal ectopias in *Eml3*-deficient embryos while determining that cortical layering outside of the focal neuronal ectopias is normal in *Eml3*-KO mice. Specifically, E.D. participated in the design and generation of *Eml3*-deficient mice, managed the mouse colonies, collected and phenotyped embryos and post-natal mice, processed tissues and performed coIP-M.S. and immunoblotting experiments, sequenced *EML3* in clinical samples. E.D. discovered the perinatal mortality of *Eml3*-KO mice and their small size, also defined EML3 protein interactions in tissues relevant to the *Eml3*-KO phenotypes.

V.P. designed and performed experiments aimed at the characterization of the onset of neurogenesis in *Eml3*-KO embryo brains.

S.B. designed and performed TIRF experiments confirming that the EML3 protein binds polymerized microtubules directly and that the EML3 protein can recruit DYNLL proteins to microtubules.

H.v.B. and M.S. provided clinical samples for sequencing the *EML3* gene.

A.B. performed molecular modeling and gave advice for the generation of DYNLL binding-deficient EML3 proteins.

S.S. provided guidance and technical support for brain phenotyping. Provided supervision and mentorship throughout the study and reviewed the manuscript.

Y.Y. provided guidance and technical support for early embryo phenotyping. Provided supervision and mentorship throughout the study and reviewed the manuscript.

R.R.M. Secured funding, participated in the design of experiments, provided supervision and mentorship throughout the study and reviewed the manuscript.

