## Supplementary material for "The microtubule-binding protein EML3 is required for mammalian embryonic growth and cerebral cortical development; *Eml3* null mice are a model of cobblestone brain malformation": Supp. Fig. 1

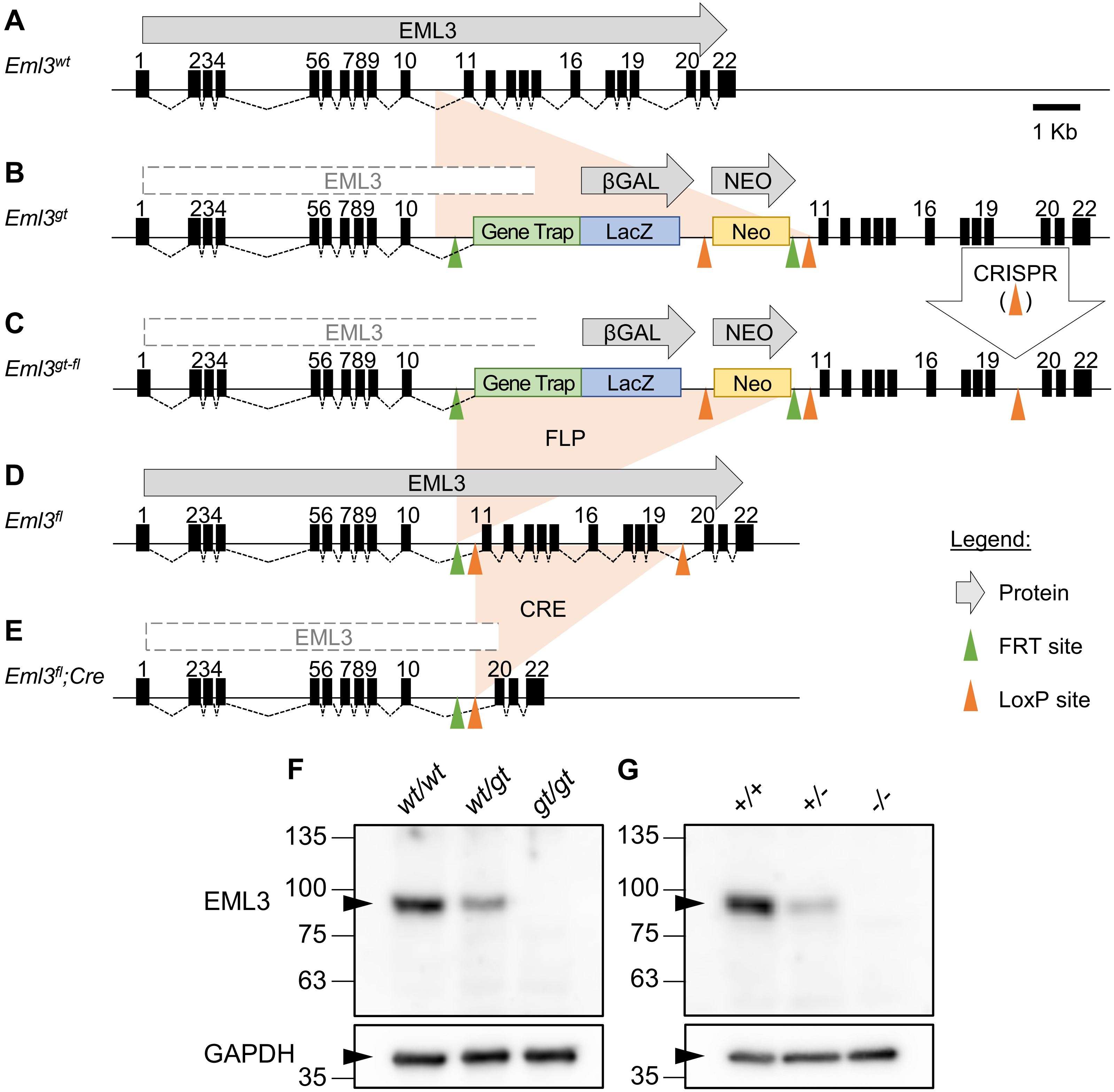

A
EML3
Eml3wt
1 234 56 789 10 11 16 19 20 22
1 Kb
B
EML3
βGAL
NEO
Eml3gt
Gene Trap
LacZ
Neo
CRISPR
C
Eml3gt-fl
FLP
D
Eml3fl
CRE
E
Eml3fl;Cre
Legend:
Protein
FRT site
LoxP site
F
wt/wt
wt/gt
gt/gt
G
+/+
+/-
-/-
135
100
75
63
35
EML3
GAPDH

**Supp. Fig. 1. Constitutive and tissue-specific targeting of *Eml3*. (A)** *Eml3* has 22 coding exons in mouse chromosome 19. **(B)** Constitutive *Eml3* null mice (*Eml3*$^{tm1e(EUCOMM)Wtsi}$ or *Eml3*$^{gt}$) with a gene trap allele from a EUCOMM ES cell line. **(C)** The gene trap-floxed allele (*Eml3*$^{em1Mci}$ or *Eml3*$^{gt\text{-}fl}$) with a LoxP site inserted into intron 19 (floxed exons 11-19). **(D)** The floxed allele (*Eml3*$^{em1.1\mathrm{Mci}}$ or *Eml3*$^{fl}$) with the gene trap cassette excised. **(E)** The tissue-specific *Eml3* null alleles, (*Eml3*$^{fl}$*;Cre*) in the presence of *Cre* recombinases under the control of tissue-specific promoters. The use of a germline-expressed *Cre* driver (*E2a-Cre*) yielded a global *Eml3* null allele named *Eml3*$^{em1.2Mci}$. **(F, G)** Immunoblotting of E15.5 whole embryo lysates with EML3 antibody (EML3-C884) detected no EML3 protein (~90kDa) in homozygotes for either **(F)** the gene trap allele (*Eml3*$^{tm1e(EUCOMM)Wtsi}$ or *Eml3*$^{gt}$), or **(G)** the exon11-19 deletion allele (*Eml3*$^{em1.2Mci}$ or *Eml3*$^{-}$). Heterozygous embryos yielded approximately half the amount of protein as wild type controls in both mouse lines. GAPDH immunoblotting was used as the protein loading control.
