## Supplementary material for "The microtubule-binding protein EML3 is required for mammalian embryonic growth and cerebral cortical development; *Eml3* null mice are a model of cobblestone brain malformation": Supp. Fig. 2

### Slide 1
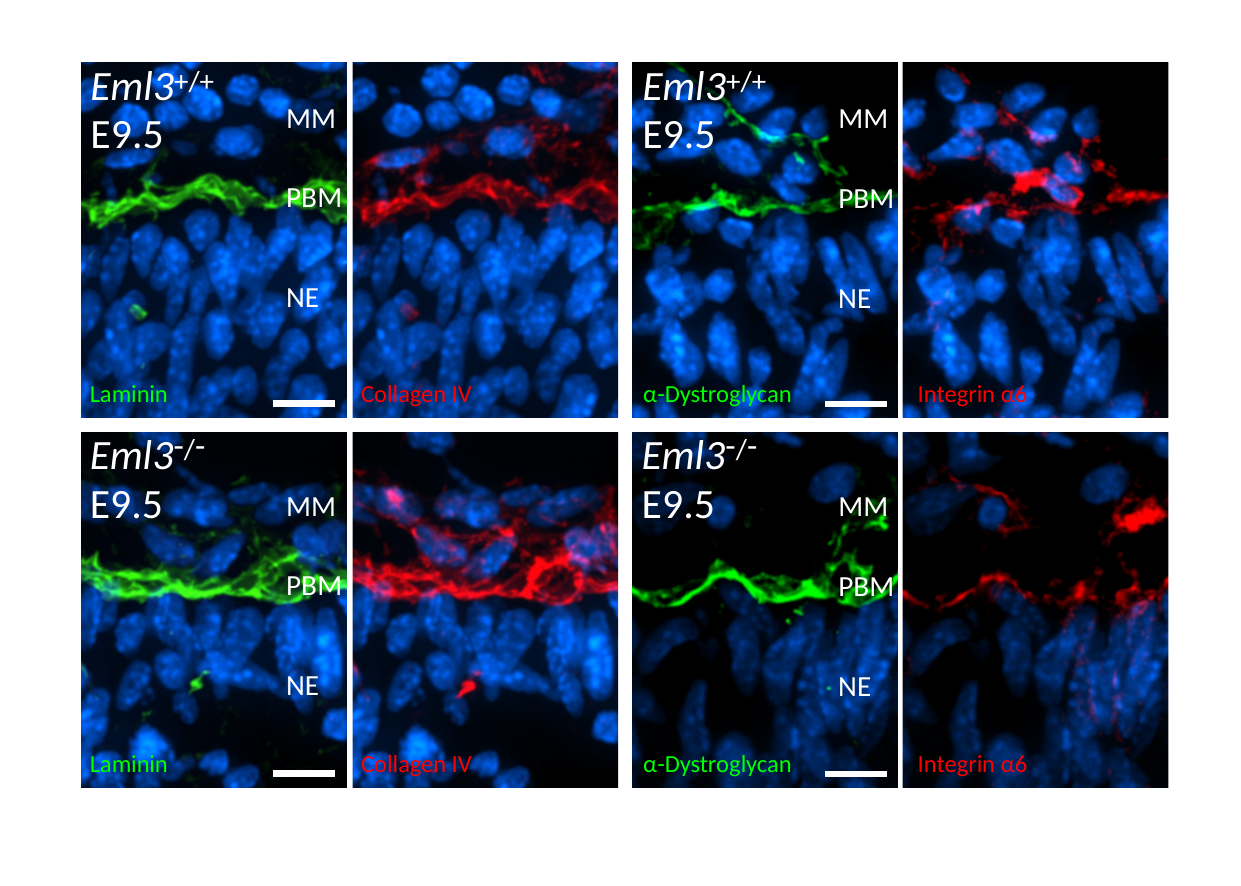

Eml3^+/+^
E9.5
Eml3^+/+^
E9.5
MM
MM
PBM
PBM
NE
NE
Laminin
Collagen IV
α-Dystroglycan
Integrin α6
Eml3^-/-^
E9.5
Eml3^-/-^
E9.5
MM
MM
PBM
PBM
NE
NE
Laminin
Collagen IV
α-Dystroglycan
Integrin α6

### Slide 2
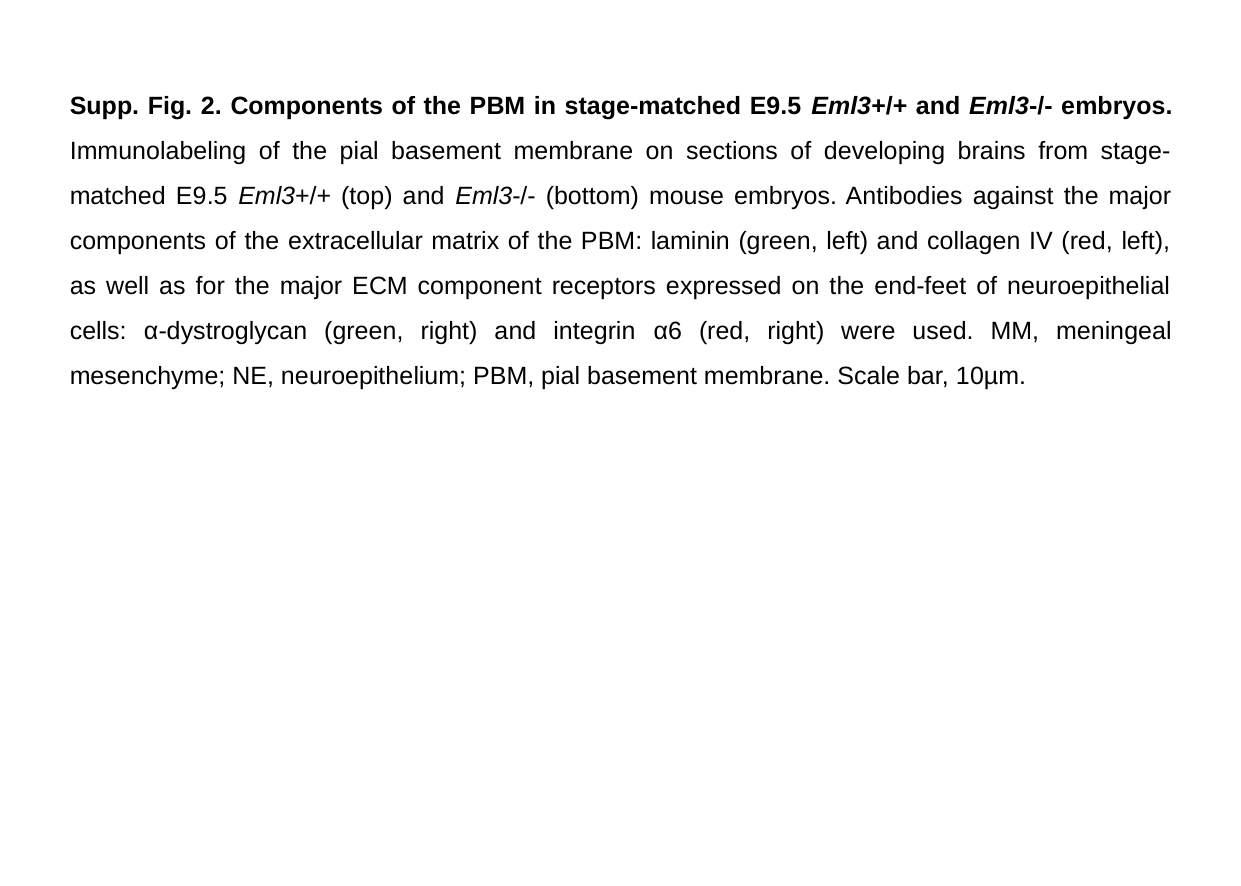

Supp. Fig. 2. Components of the PBM in stage-matched E9.5 Eml3+/+ and Eml3-/- embryos. Immunolabeling of the pial basement membrane on sections of developing brains from stage-matched E9.5 Eml3+/+ (top) and Eml3-/- (bottom) mouse embryos. Antibodies against the major components of the extracellular matrix of the PBM: laminin (green, left) and collagen IV (red, left), as well as for the major ECM component receptors expressed on the end-feet of neuroepithelial cells: α-dystroglycan (green, right) and integrin α6 (red, right) were used. MM, meningeal mesenchyme; NE, neuroepithelium; PBM, pial basement membrane. Scale bar, 10µm.
